# Two-ended recombination at a Flp-nickase-broken replication fork

**DOI:** 10.1101/2024.04.10.588130

**Authors:** Rajula Elango, Namrata Nilavar, Andrew G. Li, Erin E. Duffey, Yuning Jiang, Daniel Nguyen, Abdulkadir Abakir, Nicholas A. Willis, Jonathan Houseley, Ralph Scully

## Abstract

Collision of a replication fork with a DNA nick is thought to generate a one-ended break, fostering genomic instability. Collision of the opposing converging fork with the nick could, in principle, form a second DNA end, enabling conservative repair by homologous recombination (HR). To study mechanisms of nickase-induced HR, we developed the Flp recombinase “step arrest” nickase in mammalian cells. Flp-nickase-induced HR entails two-ended, BRCA2/RAD51-dependent short tract gene conversion (STGC), BRCA2/RAD51-independent long tract gene conversion, and discoordinated two-ended invasions. HR induced by a replication-independent break and by the Flp-nickase differ in their dependence on *BRCA1*. To determine the origin of the second DNA end during Flp-nickase-induced STGC, we blocked the opposing fork using a site-specific Tus/*Ter* replication fork barrier. Flp-nickase-induced STGC remained robust and two-ended. Thus, collision of a single replication fork with a Flp-nick can trigger two-ended HR, possibly reflecting replicative bypass of lagging strand nicks. This response may limit genomic instability during replication of a nicked DNA template.

## Introduction

Single-stranded (ss)DNA breaks (“nicks”) are common intermediates of DNA repair pathways and of the action of DNA modifying enzymes such as Type I topoisomerases^1^. Most nicks are ligated rapidly by single-stranded break repair pathways. However, if a nick persists into the S phase of the cell cycle, collision of a replication fork with the nick can convert the nick into a double strand break (DSB), a more toxic lesion that may be repaired by homologous recombination^2–4^. Some cancer chemotherapeutics act by nascent DNA strand termination, potentially flooding the replicating genome with ssDNA nicks and gaps^5^. Inhibitors of poly(ADP-ribose) polymerase (PARP), which are used to treat homologous recombination (HR)-defective cancers lacking *BRCA1* or *BRCA2*, trap PARP1 on ssDNA and delay the ligation of Okazaki fragments^6–9^. The Topoisomerase I (TopI) inhibitor and cancer therapeutic, camptothecin (CPT), acts by stabilizing a 3′-phosphotyrosyl covalent intermediate between the TopI catalytic tyrosine and the 3′ end of the DNA nick, leading to formation of a DNA-protein crosslink (DPC) and persistence of the associated nick^1^. CRISPR/Cas9 nickases are being developed for use in therapeutic gene editing and other interventions^10^. Thus, it is important to understand the mechanisms that promote or restrain genomic instability at sites of DNA nick-induced fork breakage.

Repair of a replication-independent two-ended DSB by HR in somatic mammalian cells predominantly involves the synthesis-dependent strand annealing (SDSA) pathway^11–13^. The DNA ends are engaged by BRCA1/BARD1, CtIP and the Mre11/Rad50/NBS1 (MRN) complex to initiate short range DNA end resection^14^. This process evicts the Ku heterodimer—the apical complex involved in classical non-homologous end joining (cNHEJ)—from the DNA ends and exposes ssDNA with a free 3’ end. Subsequently, Exo1, DNA2 and the Bloom’s syndrome helicase mediate long range resection extending the 3’ ssDNA tract. BRCA1 also acts downstream of resection, recruiting BRCA2/RAD51 through PALB2 and, later still, facilitating RAD51-mediated strand exchange^15–17^. BRCA2 displaces the ssDNA-binding heterotrimeric complex RPA from ssDNA to load RAD51, the central HR recombinase^18,19^. Following RAD51-mediated invasion of a homologous donor by one end of the DSB, repair synthesis initiated at the invading 3’end extends the nascent strand using the homologous donor as a template. Timely displacement of the nascent strand enables termination of gene conversion by annealing with the exposed complementary ssDNA of the homologous second end of the break, resulting in “short tract” gene conversion (STGC). STGCs of <100 bp dominate the SDSA pathway^12,20^. A minority (∼5%) of RAD51-mediated invasions trigger replicative “long tract” gene conversion (LTGC), in which several kilobases of nascent strand extension occur prior to termination^21–23^.

If the second end of the DSB lacks homology with the displaced nascent strand, termination of HR by annealing is not possible because of the lack of complementarity between the two ssDNA ends. In yeasts, a robust, mutagenic replicative mechanism termed break-induced replication (BIR) can extend nascent strand synthesis to the end of the chromosome, resulting in gene conversions of >100 kb^24,25^. In mammalian cells, the majority of HR events established in the absence of a homologous second end produce gene conversions of <1 kb, with termination mediated by end joining between the displaced nascent strand and the non-homologous second end^26^. The remaining events resolve by LTGC, which has mechanistic similarities to BIR^27^. When HR is terminated by non-homologous end joining, the extent of nascent strand synthesis (and hence the length of the gene conversion tract) varies, resulting in clone-to-clone variation in the structure of the HR repair product^26^. This variability contrasts with termination of two-ended HR by annealing, where homologous pairing ensures that HR products are of a predictable size and structure.

A long-standing model proposes that collision of the fork with a DNA nick leads to fork breakage, with the obligatory formation of a one-ended DSB^13^. Repair of such lesions is necessarily error-prone, the options including BIR-type responses or end-joining with heterologous DNA ends to form a translocation or some other chromosomal rearrangement^12,28^. In replicating frog egg extracts, fork collision with a nicked leading strand template was shown to produce one-ended breaks, caused by loss of the CMG replicative helicase from the broken leading strand along which it travels^29,30^. A similar pattern of one-ended breaks was also seen in response to a lagging strand nick. In this case, interaction of the CMG helicase with the free 5’end of the ssDNA-dsDNA boundary at the site of lagging strand breakage triggered a replication termination response, with eviction of the CMG helicase from the site by CRL2^Lrr1^ and the p97 ATPase^31,32^. One-ended DSBs are also implicated as intermediates of nickase-induced repair in yeasts, where persistent DNA nicks were found to provoke BIR at sites of fork breakage^33,34^. The extent of BIR was limited by the arrival of a second, converging replication fork from the neighboring replicon. These reports highlight the potential contribution of the opposing replication fork as a modifier of nickase-induced repair. Indeed, two-ended HR is observed at nickase-induced DSBs, but it is unresolved whether the second end arises from the collision of one fork, with replicative bypass of the nick, or from the converging fork’s collision with the nick^4^. A study in mammalian cells using Cas9 nickases revealed a bias in favor of LTGC at the site of Cas9 nickase-induced fork breakage, consistent with the presence of abundant one-ended breaks^27^.

Flp is a member of the λ integrase, or tyrosine-based family of site-specific recombinases; it catalyzes recombination between identical copies of its cognate binding site, *Frt*^35^. Each *Frt* site binds three Flp monomers, but only two of these (a and a′ in **Figure 1A**) participate directly in the recombination reaction. One of the two recombinationally active Flp monomers makes an incision *in trans* at the edge of its partner Flp binding site^36,37^. Flp-mediated incision entails attack of a 3′-phophodiester bond in one DNA strand of *Frt* by a tyrosine residue in the Flp active site, to form a covalent 3′-phosphotyrosyl DPC. Since only one of the two Flp monomers can perform this function at any given time, the solitary incision results in the formation of a DNA nick. Hydrolysis of the DPC is one step in the choreographed resolution of the site-specific Flp/*Frt* recombination reaction. The FlpH305L “step arrest” mutant is defective for this hydrolysis step^38^. As a result, the normally transient DPC is stabilized, rendering the *Frt* site nick unligatable as long as the DPC remains. In yeast, the genetics of cellular sensitivity to a FlpH305L-nick matches that of CPT, indicating that the FlpH305L-nick is a good model of the trapped, CPT-poisoned Top I-DNA cleavage complex (CPT-TopIcc)^39^. Flp introduces nicks into either DNA strand of a *Frt* site *in vitro* (incisions labeled a or a′ in **Figure 1A**)^40^. Thus, in cell-based assays, FlpH305L could potentially interrupt either the leading or lagging parental strand of a colliding replication fork.

**Figure 1.**
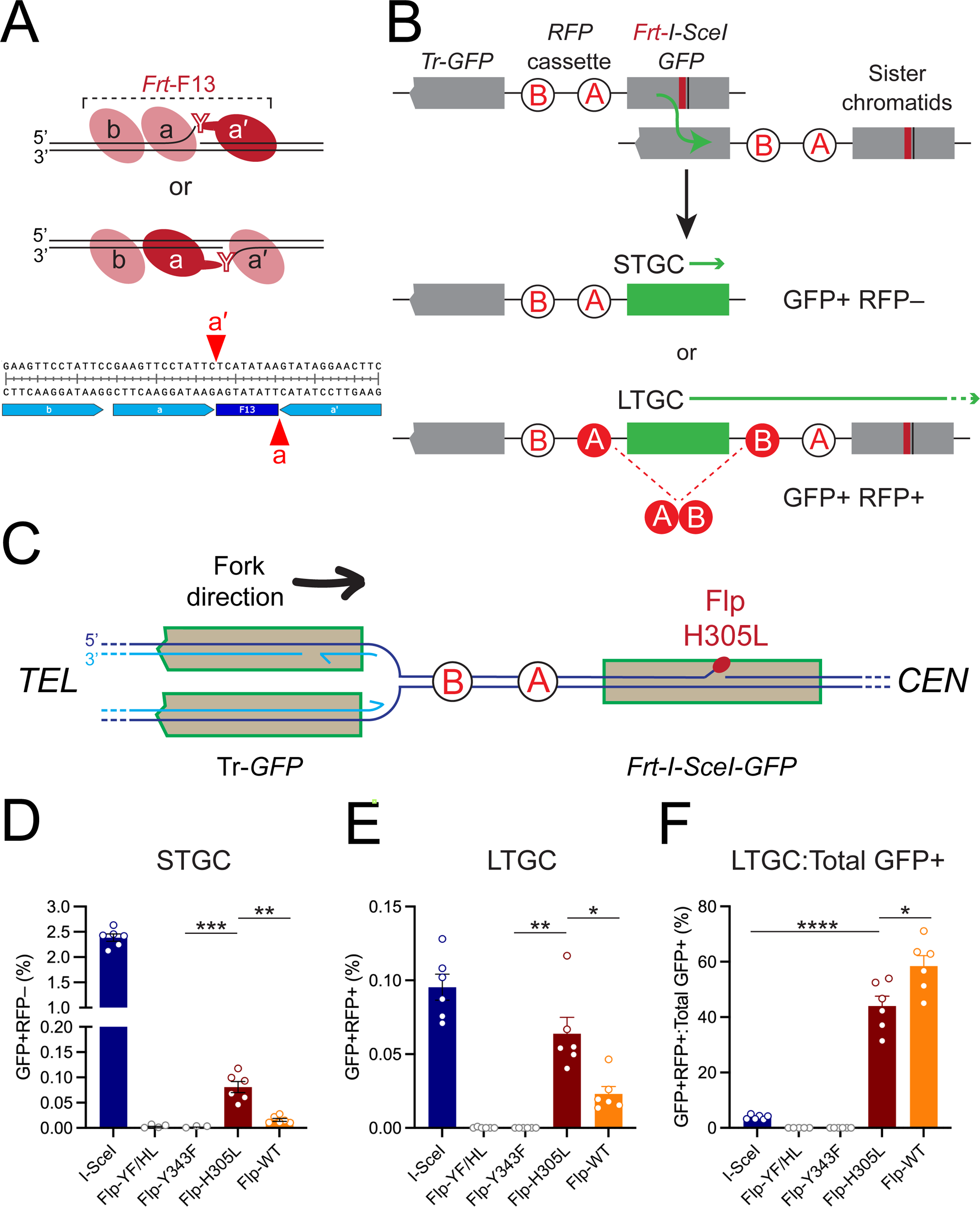
FlpH305L-induced HR products in mammalian cells. **A.** FlpH305L binding to the F13-*Frt* site generates DPC nicks on either strand. Pink: non-incising Flp monomers. Dark red: incising Flp monomer that contributes the active site tyrosine (Y, as shown). Flp monomers *a* and *a*′, but not monomer *b*, normally contribute to recombination at *Frt*. Lower panel: sequence of F13 *Frt* site in HR reporter, with sites of incisions made by Flp monomers *a* and *a*′ as shown. **B.** Schematic of HR outcomes detected by reporter. STGC generates GFP^+^RFP^−^ outcome; LTGC generates GFP^+^RFP^+^ outcome. **C.** Cartoon showing orientation of the HR reporter with respect to a replication fork traveling through *Rosa26* from *TEL* to *CEN*. **D.** STGC frequencies induced by I-SceI and Flp mutants shown. Flp-YF/HL: Y343F/H305L double mutant. Data shows mean and s.e.m. of 6 independent experiments (n=6). Unpaired *t*-test: ** P < 0.01; *** P < 0.001. **E.** and **F.** LTGC products (E) and ratio of LTGC:Total GFP^+^ frequencies (F) from the same experiments as in C (n=6). * P < 0.05; **** P < 0.0001. See also **Supplemental Figure S1**.

In this study, we develop the FlpH305L nickase system for use in mammalian cells. We find that FlpH305L-induced repair generates STGC products of two-ended HR as well as aberrant replicative repair outcomes. We define the genetic regulation of each pathway. Further, using a site-specific replication fork barrier to insulate the nick site from the converging opposing replication fork, we find that FlpH305L-induced two-ended STGC requires the arrival of only one replication fork at the nick site. This finding suggests that two-ended breaks at nickase-collapsed forks can arise from replicative bypass of the nick, without any contribution from the opposing converging fork.

## Results

### FlpH305L induces HR at a chromosomally targeted *Frt* site in mammalian cells

To study FlpH305L-induced HR in mammalian cells, we modified a previously described HR reporter^26^ by positioning a single *Frt* site within a full-length copy of *GFP*, adjacent to an 18 bp target site for the rare-cutting homing endonuclease I-SceI (**Figure 1B**). We used an artificial 8 bp spacer sequence termed F13 within the *Frt* site, so that it could not recombine with native *Frt* sites in future studies^41^. Recombination between the broken *GFP-Frt F13* heteroallele and a neighboring 5’ truncated *GFP* donor results in expression of wild type (wt)*GFP*, which is readily detected by flow cytometry. Between the two *GFP* copies we placed a previously-described *RFP* cassette, in which two artificial *RFP* exons (A and B, **Figure 1B**) are positioned non-productively^26^. Thus, STGC generates a GFP^+^RFP^−^ outcome, while LTGC expands the reporter, duplicating the *RFP* cassette and leading to expression of wt*RFP* (**Figure 1B**). LTGC products are GFP^+^RFP^+^.

To determine the major direction of DNA synthesis through *Rosa26*, we performed TrAEL-seq on unmodified mouse embryonic stem (mES) cells^42^. TrAEL-seq detects the genome-wide distribution and orientation of 3’ DNA ends, which map to nascent leading strands of replication forks^42^. TrAEL-seq analysis suggested that the majority of forks traverse *Rosa26* from telomere (*TEL*) to centromere (*CEN*), co-directional with *Rosa26* transcription (**Supplemental Figure 1A**). This conclusion is indirectly supported by our previous study using high throughput genome-wide translocation sequencing (HTGTS), in which we found that fork stalling at a Tus/*Ter* replication fork barrier (RFB) at *Rosa26* is dominated by DNA ends derived from forks traveling from *TEL*-to-*CEN*^43^. **Figure 1C** shows the orientation of a *Rosa26*-targeted *Frt*-F13 HR reporter relative to the *TEL*-to-*CEN* fork direction. We targeted a single copy of the *Frt*-F13 HR reporter to *Rosa26* in mES cells that carry a conditional allele of *Brca1*, termed *Brca1*^fl^ (see STAR Methods). Cre-mediated excision of the in-frame “floxed” exon 11 of *Brca1* generates a hypomorphic *Brca1*^Δ11^ allele, which is defective for DNA end resection but supports proliferation of mES cells^44,45^. The second *Brca1* allele, *Brca1*^hyg^, is interrupted by a hygromycin resistance cassette, and is functionally null. We identified six independent clones carrying an intact, single-copy of the *Frt*-F13 HR reporter targeted to *Rosa26*. We also generated a reporter in which the full *Frt*-F13 site was replaced by a minimal (“*Frt*-F13min”) site that retains only the two Flp binding sites that participate directly in the site-specific recombination reaction (**Supplemental Figure S1B**). We generated five independent *Brca1*^fl/hyg^ clones targeted with the *Frt*-F13min HR reporter to *Rosa26*. We constructed vectors, codon-optimized for mammalian expression, of derivatives of Flpe, a variant of Flp engineered for higher activity at 37°C^46^. These variants included wtFlp, FlpH305L (step arrest mutant), FlpY343F (catalytically inactive mutant) and the FlpH305L/Y343F double mutant. Each cDNA was fused to an in-frame N-terminal 3xmyc epitope tag to enable protein detection and a SV40 nuclear localization sequence to ensure nuclear localization (**Supplemental Figure S1C**).

Transfection of FlpH305L induced HR in the six *Frt-*F13-HR clones, yielding STGC frequencies ∼30% higher than the five F13min-HR reporter clones (**Supplemental Figure S1D**). Thus, the third Flp binding site in *Frt* (site *b* in **Figure 1A**) likely contributes to nicking efficiency or to the persistence of the FlpH305L DPC. We performed all subsequent experiments in *Frt-*F13-HR cells. No HR products were induced by the catalytically inactive mutants FlpY343F or FlpH305L/Y343F (**Figures 1D-1F** and **Supplemental Figure S1E**). This result shows that incisions at *Frt* are required for HR and that Flp-*Frt* binding *per se* does not trigger HR. Wild type Flp generated STGC frequencies above background but ∼5-fold lower than those induced by FlpH305L (**Figure 1D**). In response to an I-SceI-induced DSB, ∼95% of HR products resolve as STGC and ∼5% as LTGC in wild type cells^21,26^ (**Figure 1F**). In contrast, ∼60% FlpH305L-induced HR products resolve as STGC and ∼40% as LTGC (**Figures 1D-1F**). We also noted induction by FlpH305L of an unanticipated minor repair outcome that is GFP^−^RFP^+^ (**Supplemental Figure S1E**).

To determine the mechanisms underlying FlpH305L-induced HR, we analyzed FACS-sorted FlpH305L-induced HR products by Southern blotting. When STGC proceeds in the absence of a homologous second end, the resulting STGC products are of variable size, reflecting variation in the site of HR termination by end-joining^26^. In contrast, two-ended STGCs, in which the second end is homologous to the donor template, are of a fixed/stereotyped size that structurally resemble the parental HR reporter. Southern analysis of FACS-sorted STGC clones revealed products of a fixed, stereotyped size, indicating that they are the outcome of two-ended HR (**Supplemental Figure S1F**). FlpH305L-induced LTGCs conformed to the expected structure, similar to that reported previously for I-SceI-induced LTGC^26,47^ (**Supplemental Figure S1F**). Analysis of GFP^−^RFP^+^ products is described below.

### Classical NHEJ does not compete with Flp-nick-induced HR

Wild type Flp can introduce a DSB into a single *Frt* site *in vitro*, although the majority of products are DNA nicks^40^. The FlpH305L mutant is a less potent nuclease than wt Flp^38^, suggesting that it will act primarily as a nickase. However, it was important to determine whether the repair products induced by FlpH305L reflect its major nickase activity or whether they might be products of a minor DSB nuclease activity. In the repair of a replication-independent DSB, the cNHEJ pathway competes with HR; deletion of the core cNHEJ gene *Xrcc4* greatly elevates the frequency of DSB-induced HR^12,48^. In contrast, cNHEJ gene deletion has no impact on HR induced by a CRISPR/Cas9 nickase^49^. To determine whether cNHEJ competes with Flp-nick-induced HR, we targeted a single copy of the *Frt*-F13 HR reporter to *Rosa26* in *Xrcc4*^fl/fl^ mES cells, transduced a *Xrcc4*^fl/fl^ reporter clone with Cre recombinase adenovirus and identified clones that had deleted *Xrcc4* biallelically (*Xrcc4*^−/–^). We did not retrieve any clones in which *Xrcc4* deletion had not occurred. As expected, I-SceI-induced HR frequencies were elevated in *Xrcc4*^−/–^ reporter cells in comparison to parental *Xrcc4*^fl/fl^ cells^50^ (**Figure 2A** and **Supplemental Figure 2A**). In contrast, FlpH305L-induced HR frequencies were similar in *Xrcc4*^−/–^ and *Xrcc4*^fl/fl^ cells (**Figure 2B** and **Supplemental Figure 2B**). Thus, as was observed with CRISPR/Cas9 nickase-induced HR^49^, Flp-nick-induced HR is unaffected by the cNHEJ status of the cell. These results suggest: first, that the Flp-nick system as studied here does not generate quantitatively important frequencies of DSBs prior to fork collision; second, that the exclusion of cNHEJ from competition with HR is a general feature of nickase-induced repair pathway choice.

**Figure 2.**
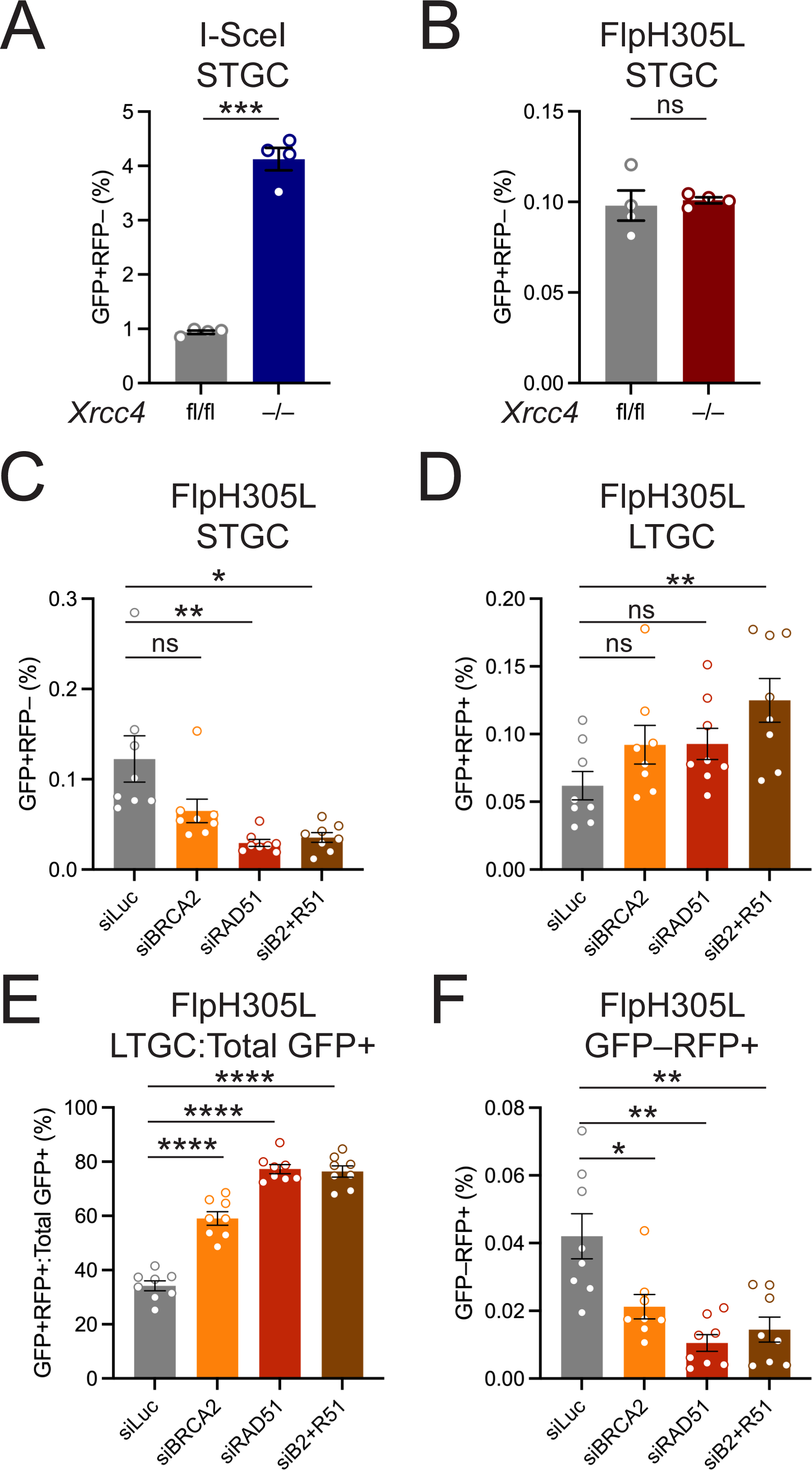
Genetic analysis of FlpH305L-induced repair. **A** and **B.** I-SceI induced (A) and FlpH305L-induced (B) STGC in parental *Xrcc4*^fl/fl^ and *Xrcc4*^−/–^ reporter cells. Data shows mean and s.e.m. of 4 independent experiments (n=4). Unpaired *t*-test: ns: not significant; *** P < 0.001. **C-F.** Cells were co-transfected with FlpH305L and siRNAs shown. See STAR Methods for experimental details. Data shows mean and s.e.m. of 8 independent experiments (n=8). Unpaired *t*-test: ns: not significant; * P < 0.05; ** P < 0.01; **** P < 0.0001. **C.** STGC frequencies. **D.** LTGC frequencies. Note: FlpH305L-induced LTGCs are independent of BRCA2 and Rad51. **E.** ratio of LTGC:Total GFP^+^ frequencies. **F.** Frequencies of FlpH305L-induced GFP^−^RFP^+^ outcome. See also **Supplemental Figure S2**.

### RAD51-dependent and -independent pathways of Flp-nick repair

To determine RAD51 dependencies of Flp-nick-induced repair, we used siRNA to deplete RAD51 and/or BRCA2 during FlpH305L-induced HR. FlpH305L-induced STGC, like I-SceI-induced STGC, was suppressed by depletion of RAD51 and/or BRCA2 (**Figure 2C** and **Supplemental Figure S2C**). In striking contrast, FlpH305L-induced LTGC was unaffected or increased in RAD51- and/or BRCA2-depleted cells (**Figure 2D**). The same siRNA treatments diminished I-SceI-induced LTGC, as expected^44^ (**Supplemental Figure S2D**). In RAD51-depleted cells, ∼80% of all FlpH305L-induced GFP^+^ products were LTGCs; the equivalent ratio in I-SceI-induced repair was ∼10% (**Figure 2E** and **Supplemental Figure S2E**). The RAD51-dependence of FlpH305L-induced STGC, taken together with the above-noted Southern analysis (**Supplemental Figure S1F**), suggests that FlpH305L-induced STGC is a product of conservative SDSA. In contrast, Flp-nick-induced LTGC is mediated by a distinct pathway independent of or suppressed by BRCA2 and RAD51. Flp-nick-induced LTGC differs mechanistically from DSB-induced LTGC, since the latter is RAD51-dependent^44^ (**Supplemental Figure S2D**).

We found that FlpH305L-induced GFP^−^RFP^+^ repair products are dependent on BRCA2 and RAD51, similar to FlpH305L-induced STGC (**Figure 2F**; compare with **Figure 2C**). Quantitatively, ∼25% of all RAD51-mediated HR events at FlpH305L resolve as the GFP^−^RFP^+^ outcome. I-SceI-induced GFP^−^RFP^+^ repair products are also RAD51-dependent (**Supplemental Figure S2F**). However, they account for a much smaller fraction (∼2%) of all HR products.

### Analysis of FlpH305L-induced GFP^−^RFP^+^ events

To identify mechanisms underlying the FlpH305L-induced GFP^−^RFP^+^ outcome, we FACS-sorted GFP^−^RFP^+^ clones and analyzed their structure by Southern blotting. Six of eight FlpH305L-induced GFP^−^RFP^+^ clones were of identical structure, resembling LTGC products in size and containing three copies of *GFP*. However, unlike LTGC, in which the central *GFP* copy is wild type and only the third *GFP* copy contains an *I-SceI* site, GFP^−^RFP^+^ products contained two copies of *Frt-I-SceI-GFP* but no wild type *GFP* copy (**Figures 3A** and **3B**). Two clones (lanes 3 and 4, **Figure 3A**) revealed a different structure that likely contains three copies of *Frt-I-SceI-GFP*, as suggested by the increased intensity of the 3.5 kb restriction product with concomitant loss of the 3.8 kb band. Seven of nine I-SceI-induced GFP^−^RFP^+^ products revealed the same structure as the major FlpH305L-induced outcome (**Supplemental Figure S3A**). The remaining two I-SceI-induced GFP^−^RFP^+^ products lacked any *I-SceI* sites and their provenance is unclear. Mechanistically, each copy of *Frt-I-SceI-GFP* in the major GFP^−^RFP^+^ product, depicted in **Figure 3B**, must arise by recombination with the intact *Frt-I-SceI-GFP* copy of the sister chromatid. If so, the GFP^−^RFP^+^ product is the outcome of two independent sister chromatid recombination events. One way that this could arise is *via* discoordinated, RAD51-mediated invasions by each end of the nickase-induced break, followed by long tract gene conversion, with termination by annealing (**Figure 3C**; see **Supplemental Figure S3B** for a more detailed model). Alternative mechanisms, including rolling circle amplification of the *Frt-I-SceI-GFP* sequence, might explain the origin of the two FlpH305L-induced GFP^−^RFP^+^ repair products that contain three copies of *Frt-I-SceI-GFP*.

**Figure 3.**
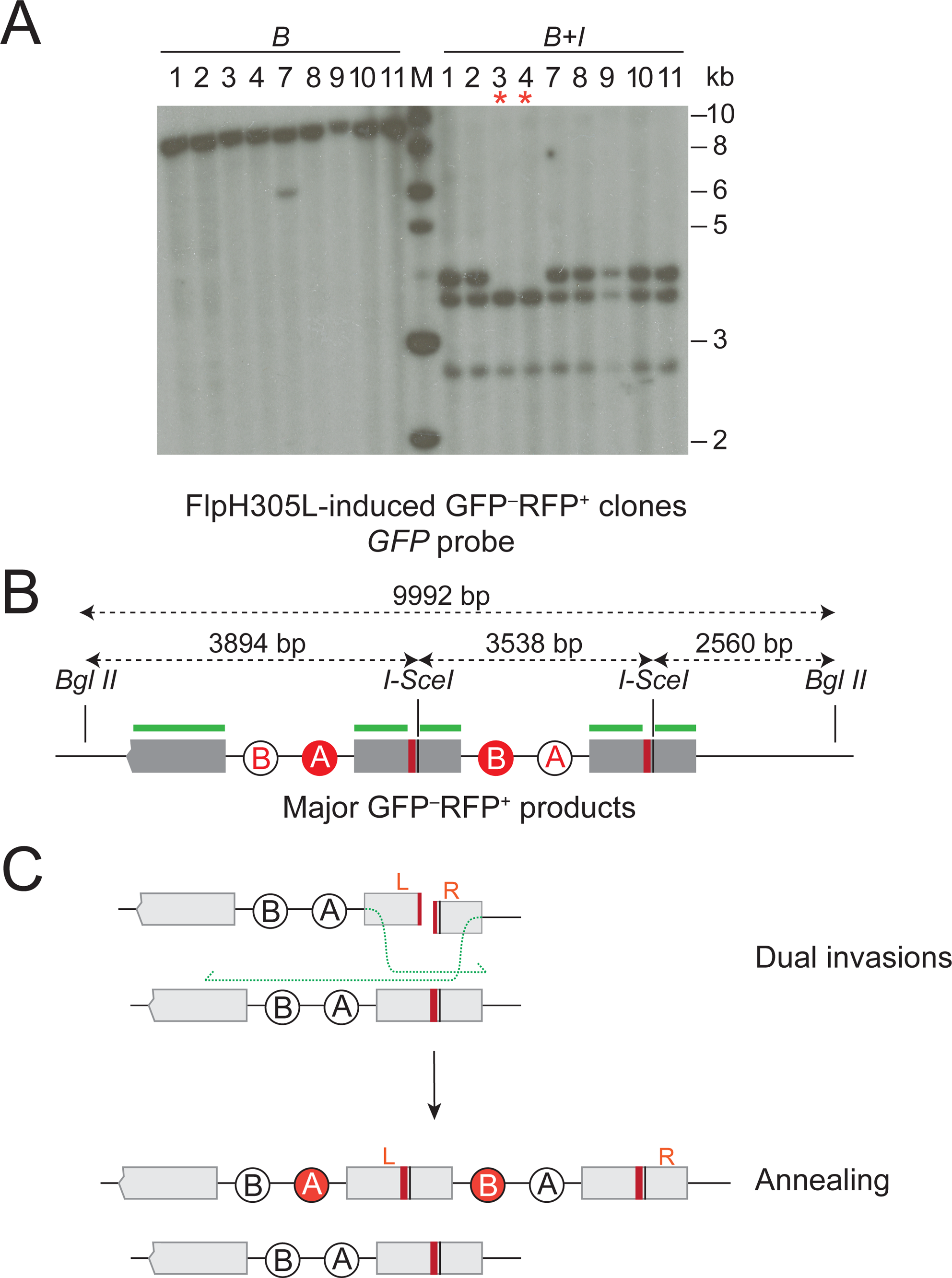
Analysis of FlpH305L-induced GFP^−^RFP^+^ products. **A.** Southern blot analysis of 9 independent FlpH305L-induced GFP^−^RFP^+^ products (clones #5 and 6 were lost). Genomic DNA was digested *in vitro* with BglII (B) or with BglII+I-SceI (B+I), *GFP* probe. Note: all clones except #3 and #4 (red *) contain two I-SceI sites. Clones #3 and #4 may contain three I-SceI sites, generating double intensity band migrating at 3538 bp. **B.** Deduced structure of major FlpH305L-induced GFP^−^RFP^+^ products, with predicted restriction fragment sizes shown. Green lines: *GFP* sequences recognized by the *GFP* probe. **C.** Hypothetical mechanism of generation of FlpH305L-induced GFP^−^RFP^+^ outcome *via* discoordinated dual invasions of the sister chromatid. Original sites of left and right DNA ends are marked L and R, respectively. See also **Supplemental Figure S3**.

Note that the Flp-nick- and DSB-induced GFP^−^RFP^+^ products reported here are structurally distinct from the GFP^−^RFP^+^ non-homologous tandem duplications (TDs) observed in response to a Tus/*Ter*-induced replication fork barrier (RFB) in *Brca1* mutant cells—a model of TDs that arise in human *BRCA1*-mutant breast and ovarian cancers^43,51^. They are also mechanistically distinct, since TDs form by a RAD51-independent mechanism^43^. Our findings suggest that *BRCA1*-linked TDs originate from replication fork stalling events but not from nick-induced replication fork breakage.

### BRCA1-mediated DNA end resection is dispensable for Flp-nick-induced STGC

To study the contribution of BRCA1 to FlpH305L-induced HR, we used Cre transduction to generate 6 derivative clones of *Frt*-F13-HR reporter cells that were *Brca1*^Δ11/hyg^ (DNA end resection-defective hypomorph lacking *Brca1* exon 11) and 6 that were *Brca1*^fl/hyg^ (functionally wild type) (**Supplemental Figure 4A**). Note that each clone tested in these experiments had been exposed to adeno-Cre and thus the *Brca1*^Δ11/hyg^ and *Brca1*^fl/hyg^ clones studied here are truly isogenic. As reported previously, *Brca1*^Δ11/hyg^ cells revealed a reduction in I-SceI-induced STGC and a modest bias in favor of LTGC^26^ (**Figures 4A**, **4B** and **4C** and **Supplemental Figures 4B**, **4C** and **4D**). Strikingly, *Brca1*^Δ11/hyg^ and *Brca1*^fl/hyg^ *Frt*-F13-HR cells revealed identical frequencies of FlpH305L-induced STGC, suggesting that BRCA1’s function in DNA end resection is dispensable for FlpH305L-induced STGC (**Figure 4A** and **Supplemental Figure 4B**). FlpH305L-induced LTGC was increased in *Brca1*^Δ11/hyg^ F13-HR cells in comparison to isogenic *Brca1*^fl/hyg^ cells (**Figures 4B** and **4C**; **Supplemental Figures 4C** and **4D**), indicating that BRCA1 suppresses LTGC at Flp-nick-broken forks. We noted a small but statistically significant increase in FlpH305L-induced GFP^−^RFP^+^ products in *Brca1*^Δ11/hyg^ cells in comparison to *Brca1*^fl/hyg^ cells, and a similarly modest reduction in the equivalent I-SceI-induced product (**Figure 4D** and **Supplemental Figure 4E**).

**Figure 4.**
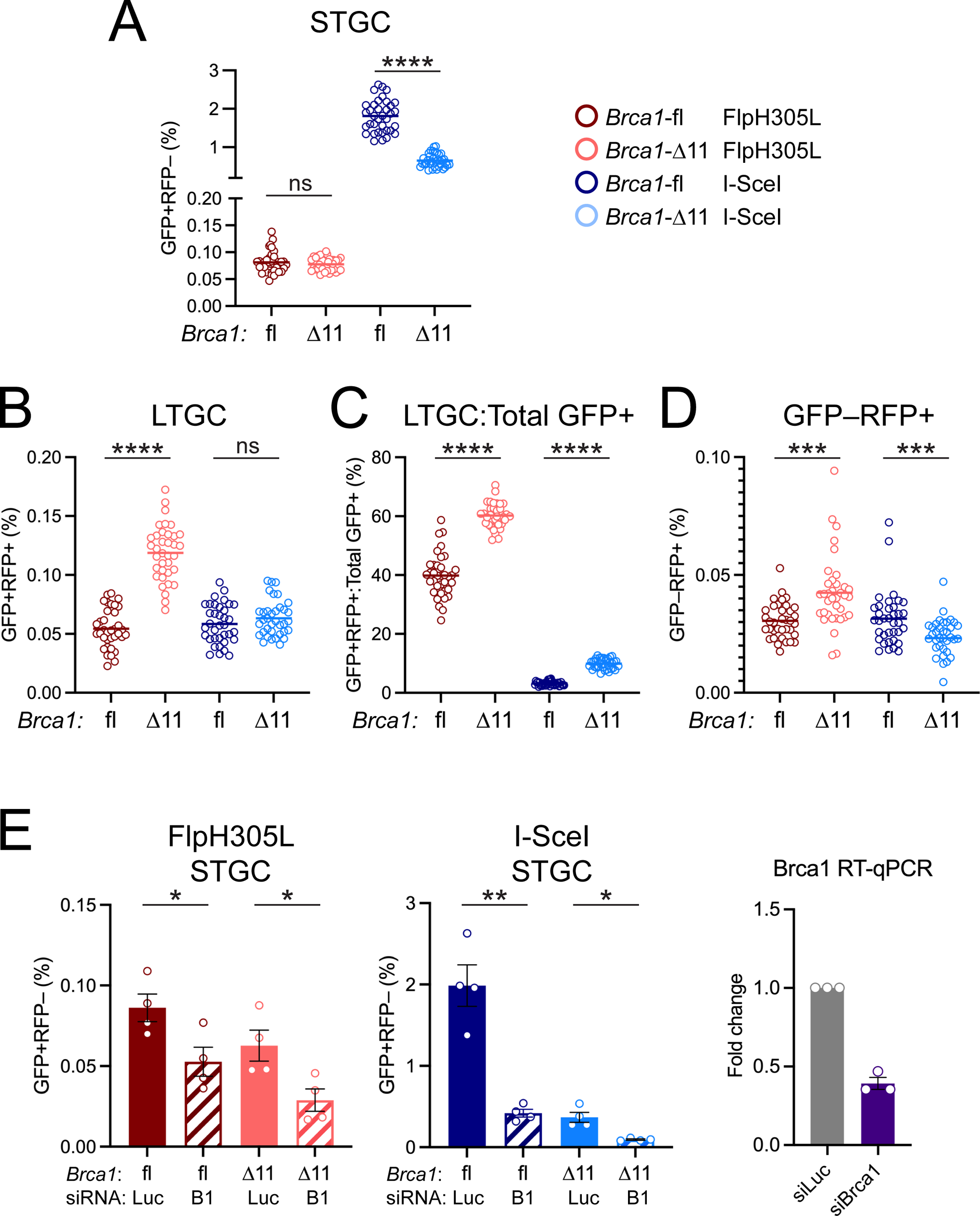
*Brca1* exon 11 is dispensable for FlpH305L-induced STGC. Data shown is pooled from 6 independent Cre-exposed Brca1^fl/hyg^ clones and 6 independent Cre-exposed Brca1^Δ11/hyg^ clones. See **Supplemental Figure S4** for data corresponding to individual clones in the same experiments. Data shows mean and s.e.m. of 6 independent experiments (n=6). Unpaired *t*-test: ns: not significant; * P < 0.05; ** P < 0.01; *** P < 0.001; **** P < 0.0001. **A.** STGC products. Key to right of figure shows colors corresponding to specific combinations of *Brca1* genotype and transfection with FlpH305L or I-SceI. Note deletion of *Brca1* exon 11 has no impact on FlpH305L-induced STGC, but reduces I-SceI-induced STGC. **B.** LTGC frequencies. Note elevated FlpH305L-induced LTGC frequencies in Brca1^Δ11/hyg^ cells. **C.** Ratio of LTGC:Total GFP^+^ frequencies. **D.** Frequencies of FlpH305L-induced GFP^−^RFP^+^ outcomes. **E.** Brca1^fl/hyg^ clone #12 and Brca1^Δ11/hyg^ clone #8 (see **Supplemental Figure S4**) were co-transfected with siRNAs shown together with either FlpH305L or I-SceI, as described in STAR methods. Data shows mean and s.e.m. of STGC values from 4 independent experiments (n=4), with analysis by unpaired *t*-test. See also **Supplemental Figures S4** and **S5**. Right panel: RT-qPCR for *Brca1* mRNA in cells transfected with the siRNAs shown.

To determine whether regions of BRCA1 other than those encoded by exon 11 regulate FlpH305L-induced HR, we used siRNA to deplete BRCA1 in *Brca1*^Δ11/hyg^ and *Brca1*^fl/hyg^ *Frt*-F13-HR reporter cells. Depletion of BRCA1 reduced FlpH305L-induced and I-SceI-induced STGC in both *Brca1*^Δ11/hyg^ and *Brca1*^fl/hyg^ cells (**Figure 4E**). Therefore, regions of BRCA1 other than those encoded by exon 11 participate in FlpH305L-induced STGC. FlpH305L-induced LTGC frequencies were unaffected by BRCA1 depletion, leading to a modest bias in favor of LTGC over the STGC outcome (**Supplemental Figures 5A** and **5B**). I-SceI-induced LTGC followed the same pattern of dependencies that we reported previously^26,44^ (**Supplemental Figures 5A** and **5B**). FlpH305L-induced GFP^−^RFP^+^ frequencies were unaffected by BRCA1 depletion, while I-SceI-induced GFP^−^RFP^+^ frequencies were reduced by BRCA1 depletion in *Brca1*^fl/hyg^ but not *Brca1*^Δ11/hyg^ cells (**Supplemental Figure 5C**).

### Two-ended STGCs following encounter of a single replication fork with the Flp-nick

The generation of two-ended HR intermediates at FlpH305L-broken forks is at odds with the one-ended break model of nickase-induced fork breakage^13,29^. However, a second DNA end could be produced if the opposing, converging fork from the neighboring replicon were to collide with the reconstituted Flp-nick^4,33,34^. If so, blocking access of the second fork to the nick site should suppress FlpH305L-induced STGC or, at a minimum, convert the underlying repair events to one-ended HR reactions (**Figure 5A**). We therefore constructed a new reporter in which an array of six *Ter* repeats is placed ∼750 bp centromeric to the *Frt*-F13 HR reporter elements, positioned so that a Tus/*Ter* RFB prevents arrival of the second opposing leftward fork in **Figure 5A**^44^. We targeted this new reporter as a single copy to *Rosa26* and confirmed the function of the Tus/*Ter* RFB in two ways. First, to confirm that Tus binds the Ter array, we used anti-HA CUT&RUN-qPCR for HA-tagged Tus^52,53^, detecting ∼60-fold enrichment of Tus at the *Ter* array in Tus-transfected cells in comparison to empty-vector-transfected controls (**Supplemental Figure S6A**). Second, we made use of the fact that RAD51 is an early responder to the Tus/*Ter*-stalled fork^50,54^. Using anti-RAD51 CUT&RUN-qPCR, we detected ∼12-fold enrichment of RAD51 at the *Ter* array in Tus-transfected cells in comparison to EV-transfected cells—a stronger enrichment than is observed at an I-SceI-induced DSB (**Supplemental Figure S6B**). We studied the impact of the Tus/*Ter* RFB on FlpH305L-induced HR in five independent *Frt*-F13 6x*Ter* HR reporter clones. Notably, co-expression of Tus with FlpH305L had no effect on the efficiency of FlpH305L-induced STGC (**Figure 5B** and **Supplemental Figure S6C**). We noted a consistent reduction in FlpH305L-induced LTGC when the Tus/*Ter* RFB was active, but no effect on FlpH305L-induced GFP^−^RFP^+^ frequencies (**Figures 5C**, **5D** and **5E** and **Supplemental Figures S6D**, **S6E** and **S6F**). We FACS sorted FlpH305L-induced STGC clones from cells in which the Tus/*Ter* RFB was active and analyzed their structure by Southern blotting. All FlpH305L-induced STGCs in Tus/*Ter* RFB-active cells revealed repair products of a stereotyped size that co-migrated with the parental reporter (**Figure 5F**). This finding indicates that FlpH305L-induced STGC in Tus/*Ter* RFB-active cells is a product of two-ended HR. Southern analysis of FlpH305L-induced LTGCs and GFP^−^RFP^+^ clones in which the Tus/*Ter* RFB was active revealed the same structures as repair products observed in the original *Frt*-F13 reporter cells (**Supplemental Figures S6G** and **S6H**). We conclude that the arrival of a single fork at a FlpH305L-induced nick can give rise to a two-ended break, leading to repair by SDSA.

**Figure 5.**
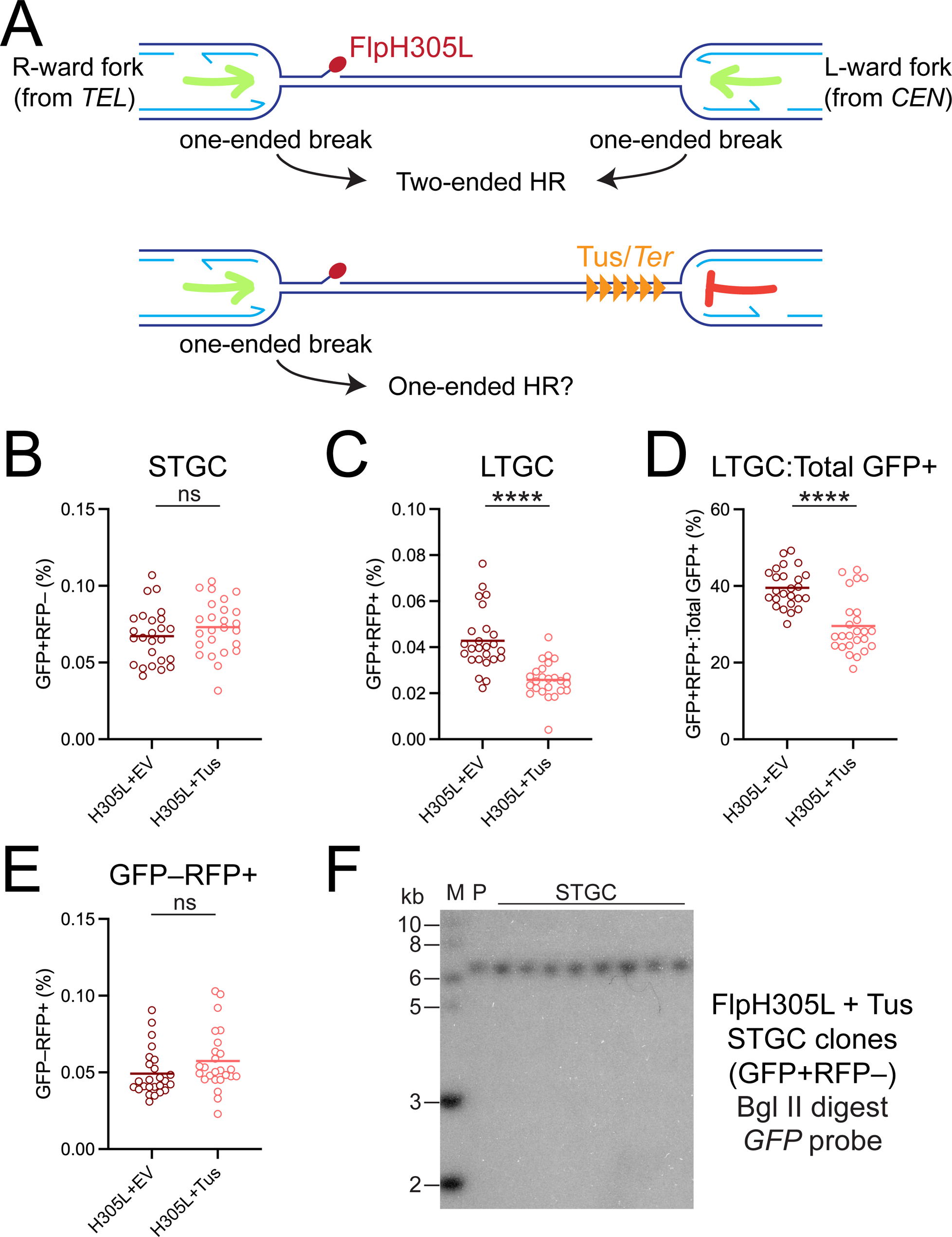
FlpH305L-induced STGC is the product of a single fork collision with the Flp-nick. **A.** Upper panel: A two-ended DSB might form from two one-ended breaks at the Flp-nick site. Collision of the rightward fork with the nick generates one DNA end. After reconstitution of the nick, collision of the opposing leftward fork from the neighboring replicon with the nick generates a second DNA end. Green arrows: direction of forks approaching Flp-nick DPC. Lower panel: A Tus/*Ter* replication fork barrier (RFB) will prevent arrival of the leftward fork, potentially blocking formation of a two-ended break at the Flp-nick site and generating only one-ended recombination products. **B.**-**E.** Data shown is pooled from 5 independent *Brca1*^fl/hyg^ clones targeted with the F13*Frt*-HR reporter carrying an additional 6x*Ter* array positioned to establish a Tus-mediated RFB to block the leftward fork. See **Supplemental Figure S6** for data corresponding to individual clones in the same experiments. Data shows mean and s.e.m. of 5 independent experiments (n=5). Unpaired *t*-test: ns: not significant; **** P < 0.0001. **B.** FlpH305L-induced STGC frequencies are unaffected by activation of the Tus/*Ter* RFB. **C.** and **D.** FlpH305L-induced LTGC frequencies (C), and the ratio of LTGC:Total GFP^+^ (D) are reduced by activation of the Tus/*Ter* RFB. **E.** FlpH305L-induced GFP^−^RFP^+^ frequencies are unaffected by activation of the Tus/*Ter* RFB. **F.** Southern blot analysis of FlpH305L-induced STGC products generated in presence of the Tus/*Ter* RFB. gDNA digested with BglII; see **Supplemental Figure S1F** for expected fragment sizes. M: MW markers. P: parental reporter cells. Note that all FlpH305L-induced STGCs are the product of two-ended HR, even in the presence of the Tus/*Ter* RFB. See also **Supplemental Figure S6**.

## Discussion

Experiments described here reveal distinct and new classes of repair products that arise at a defined DPC-site-specific nick induced by a Flp step arrest mutant in mammalian cells. One feature of the Flp-nick system is its ability to trigger two-ended STGC, reflecting the engagement of conservative SDSA. Our finding that the arrival of one replication fork at the Flp-nick site can trigger two-ended STGC challenges the long-standing model that a single fork collision with a DNA nick necessarily generates a one-ended break. This distinction is of considerable importance, since one-ended breaks may be difficult to repair by error-free mechanisms, and are likely instigators of genomic instability^12^. In contrast, as shown here, two-ended breaks that form at nickase-broken forks can be repaired in a conservative or error-free manner. The two-ended HR mechanism revealed here is likely important as a barrier to genomic instability during replication across a nicked DNA template.

A previous report of two-ended HR at a nickase-induced break considered two potential hypotheses to explain this phenomenon^4^. First, the collision of each converging fork with the nick could generate two independent one-ended breaks that are then repaired by HR as if a two-ended break (**Figure 5A**). Second, two-ended breaks might arise directly from the collision of one replisome with a lagging strand nick, followed by replicative bypass of the lagging strand nick (**Figure 6**). This model is grounded in the fact that the CMG helicase translocates along the leading parental strand^30^, making it possible, in principle, for CMG to remain topologically associated with the intact leading strand during lagging strand nick bypass. Our data excludes the first hypothesis, but is consistent with the lagging strand nick bypass model. The lack of strand specificity of the Flp-nick system limits our ability to test directly whether bypass is restricted to a lagging strand nick. However, several lines of evidence support this idea. First, a leading strand nick is predicted to result in direct loss of the CMG helicase from DNA and seems to present no opportunity for replisome bypass (**Supplemental Figure S7**). Second, Strumberg *et al*. found that conversion of the CPT-TopIcc (which the Flp-nick system models^39^) into replication-mediated DSBs in mammalian cells was detectable for leading strand nicks but not for lagging strand nicks^55^. A possible explanation of this strand asymmetry is that replication-associated leading strand nicks generate one-ended breaks that are repaired slowly, whereas lagging strand nicks generate two-ended breaks that are rapidly repaired by sister chromatid recombination/SDSA. Third, recent work from Nussenzweig and co-workers shows that CMG can bypass a lagging strand nick generated by the RuvC domain-defective Cas9 D10A nickase (personal communication; manuscript under review).

**Figure 6.**
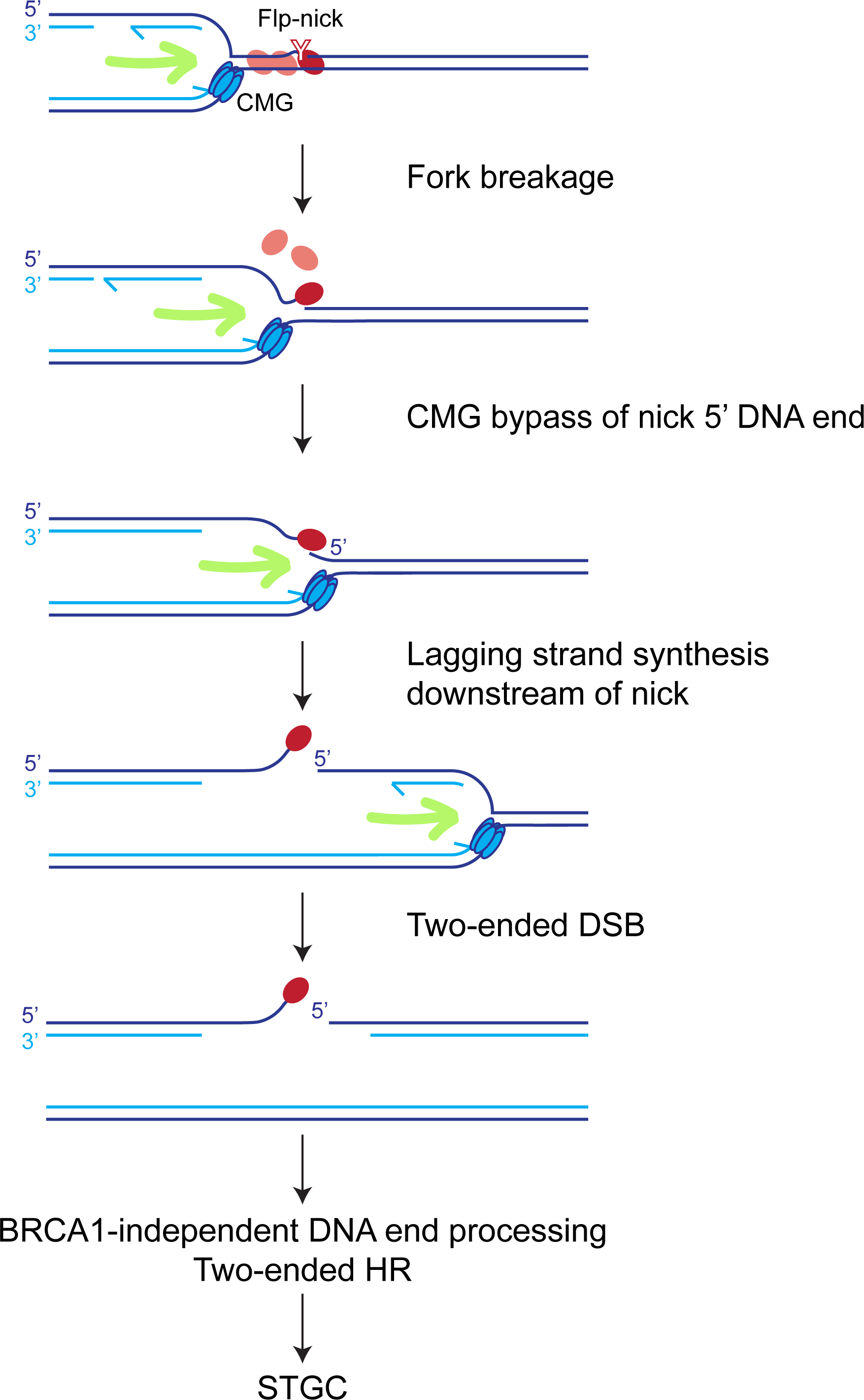
Model: collision of a single replication fork with a lagging strand Flp-nick generates a two-ended break. The replisome—traveling rightward in this cartoon—bypasses a lagging strand Flp-nick, allowing replication to continue beyond the nick site and leaving in its wake a two-ended break, generated with no contribution from the opposing (leftward) replication fork. Non-incising Flp monomers shown in pink. Incising Flp monomer shown in dark red, with active site tyrosine (Y) shown. CMG replicative helicase is shown on the uninterrupted leading strand. Green arrow shows direction of replication fork. See also **Supplemental Figure S7**.

A critical determinant of the ability of the CMG helicase to bypass a lagging strand nick may be the status of the ssDNA/dsDNA boundary formed by the 5’ DNA end of the lagging strand nick and the intact leading parental strand (**Figure 6**). The CMG helicase translocates over intact ssDNA-dsDNA junctions at the leading edge of DNA duplexes in ssDNA, without unwinding the duplex^56,57^. However, this transition in CMG-DNA association, in which the parental lagging strand is no longer excluded, acts as a replication termination signal, leading to CMG displacement from chromatin by the ubiquitin p97 pathway^31,32^. This was the case in the frog egg extract nick system of Vrtis *et al*., where CMG displacement following encounter with a lagging strand nick led to the formation of a one-ended lagging strand break^29,31,32^. However, if the 5’ end of the nick were already separated from the parental leading strand at the time of collision of CMG with the nick site, the CMG might continue to exclude the displaced lagging strand and could then bypass the nick, with resumption of lagging strand synthesis beyond the nick site (**Figure 6**). In the context of a Cas9 D10A lagging strand nick, the sgRNA target strand is nicked, leaving the complementary displaced leading strand template intact. The 5’ end of the nick is therefore masked by Cas9/sgRNA binding and separated from the leading strand at the time of fork collision, establishing the conditions for replicative bypass of the nick. In the case of the Flp nickase or the CPT-TopIcc, the free 5’ end of the nick might also be separated from the complementary leading template strand. Flp binding distorts the *Frt* site, and strand separation of the 5’ end of the Flp-nick is a scheduled step in the full recombination reaction promoted by Flp^37^. TopI has an extended footprint on both sides of the nick and, in the context of the CPT-TopIcc, CPT itself interacts with and displaces the +1 guanine base at the 5’ end of the nick^58,59^. Thus, both Flp-nick and CPT-TopIcc may, by mechanisms quite different from those of a Cas9 nickase, favor displacement of the nick 5’ end, promoting CMG bypass of the lagging strand nick.

In Vrtis *et al*., no nickase protein was present at the nick site (although it is possible that the nick provoked PARP activation at the nick). In contrast, the Flp-nick, CPT-TopIcc and Cas9 nickases each present a barrier to the replication fork at the immediate site of the nick. Even if they are weak RFBs that are ultimately displaceable, these barriers might cause transient pausing of the CMG helicase and, potentially, fork reversal. Indeed, the CPT-TopIcc is known to provoke fork reversal^60–62^. Subsequent negotiations between the replisome, the nickase complex and the fork reversal machinery may gain time for displacement of the 5’ end of the nick. In this regard, the frog egg extract nick system may resemble a conventional replication termination event, whereas the Flp-nick, CPT-TopIcc or Cas9-nickase systems may not.

STGC induced by a Flp-nick-induced DSB and by a replication-independent DSB have many similarities, including their RAD51-dependence and the structure of the final repair product. However, the two pathways diverge in regard to their dependency on BRCA1. BRCA1 exon 11 plays a key role in DNA end resection at a replication-independent DSB^12,45^, but is dispensable for Flp-nick-induced STGC (**Figure 4A**). Regions of BRCA1 other than exon 11 are, however, implicated in Flp-nick-induced STGC (**Figure 4E**). It remains to be determined which BRCA1 domains mediate this function. Our findings suggest that the regulation of DNA end resection differs between a Flp-nick-induced DSB and a replication-independent DSB. We propose that the immediate environment of the broken fork is heavily biased towards the activation of DNA end resection, to the extent that BRCA1’s normally decisive role in enabling resection at a replication-independent DSB is dispensable for resection of the nick-induced DSB. Interestingly, the cNHEJ pathway, which competes with conventional DSB-induced HR, does not compete with HR induced by either a Cas9 nickase^49^ or a Flp-nickase (**Figure 2B**). This finding might also be explained by the existence of a specialized “pro-resection” environment at the broken fork. Rapid DNA end resection could displace Ku from DNA ends, thereby aborting cNHEJ and rendering the cNHEJ status of the cell irrelevant to subsequent repair pathway choices. Similar reasoning might explain why two-ended STGC induced by Tus/*Ter* RFB-stalled forks is unaffected by the cell’s cNHEJ status^50^. In this regard, the critical DNA end processing complex MRN localizes to sites of replication in mammalian cells^63^; perhaps it is poised for immediate action at DNA ends if the fork stalls or collapses.

We find that a Flp-nick induces a higher proportion of LTGC products than does a site-specific DSB. This may indicate that a larger fraction of Flp-nick-induced recombination intermediates are one-ended, as might be expected from collisions of the replisome with leading strand Flp-nicks (**Supplemental Figure S7**). In contrast to DSB-induced LTGC, we find that Flp-nick-induced LTGC is either independent of or suppressed by BRCA1, BRCA2 and RAD51. LTGC induced at a Tus/*Ter* RFB behaves similarly^44^. In this regard, LTGC at stalled or broken forks may resemble BIR-mediated alternative lengthening of telomeres, which is also RAD51-independent^64,65^. A previous study of nickase-induced recombination, in which the HR donor molecule was episomal, reported that recombination is BRCA2- and RAD51-dependent if the donor DNA is double-stranded, but is suppressed by BRCA2 and RAD51 if the donor DNA is nicked or single-stranded^66^. Perhaps a common feature of these RAD51-independent BIR-type responses is a gapped donor template. It is interesting that activation of a Tus/*Ter* RFB near the Flp-nick site suppresses Flp-nick-induced LTGC, suggesting a possible role for the opposing fork in forming the LTGC outcome.

We found that a Flp-nick induces a previously unreported class of aberrant RAD51-dependent GFP^−^RFP^+^ repair outcomes, which is also a minor feature of I-SceI-induced repair. Our analysis of Flp-nick-induced GFP^−^RFP^+^ repair products shows that they are the product of two-ended HR and that they do not require the arrival of the second fork. These repair products therefore appear to be an alternative outcome of two-ended recombination arising from the collision of one fork with the Flp-nick. We have suggested that the major GFP^−^RFP^+^ products arise from discoordinated invasions of the sister chromatid, with resolution of the repair process by annealing. However, we cannot exclude a role for rolling circle-type repair in some of the GFP^−^ RFP^+^ outcomes. Even in this latter case, the stereotyped size of all GFP^−^RFP^+^ repair products, as revealed by Southern blotting, suggests that all GFP^−^RFP^+^ repair events are products of two-ended HR. Discoordinated invasions of this type might be a source of genomic instability, since some intermediates of this process might resolve not by local annealing, but by translocation. Notably, none of the GFP^−^RFP^+^ products studied here, either in response to a Flp-nick or a conventional DSB, were non-homologous tandem duplications (TDs) of the type observed in the response to a Tus/*Ter* RFB and in the genome of *BRCA1*-linked breast and ovarian cancers^43^. This finding suggests that *BRCA1*-linked cancer-associated TDs arise from collision of replication forks with naturally-arising replication fork barriers, but not from collisions with DNA nicks.

In summary, we find that a major product of DPC-nick-induced fork breakage is two-ended, error-free STGC and that this pathway requires the arrival of only one replication fork at the nick site. This error-free pathway may be a bulwark against genomic instability during replication of a nicked DNA template. If this observation were to extend to other nickase systems, as appears to be the case for the CRISPR/Cas9 nickase, it may have therapeutic relevance. Specifically, gene-editing nickases that target the parental lagging strand may carry a lower risk of genomic instability than those that target the parental leading strand. It will be important to test this hypothesis as these new therapeutic tools progress towards use in the clinic.

## Acknowledgements

We thank Drs. Andre Nussenzweig and Lorraine Symington for sharing unpublished data during preparation of this manuscript, and Drs. Yves Pommier, Johannes Walter and Phoebe Rice for helpful discussions. This work was supported by grants R35CA263813 to R.S., BBSRC BI Epigenetics ISP BBS/E/B/000C0523 to J.H., and Wellcome Investigator award 210754/Z/18/Z to A.A.

## Author Contributions

R.E, N.N, A.G.L., E.E.D, Y.J., D.N., A.A. and N.A.W. performed the experiments. R.E., J.H. and R.S. designed the experiments and wrote the paper.

## STAR Methods

### Molecular biology and siRNAs

The Frt-F13 and Frt-F13min - HR reporters used were assembled using standard cloning methods, similar to what was previously described for the 6x*Ter*-HR reporter **bb**. The Frt-F13-6x*Ter* reporter was generated by cloning 6x*Ter* into the AscI site, approximately 600 bp from the Frt-F13 site. wtFlp, FlpH305L, FlpY343, FlpH305L/Y343F double mutant, Tus and I-SceI plasmids were cloned into a pcDNA3β-Myc-NLS derivative vector as described in **bb**. *Ter*-containing plasmids were amplified in JJC33 (Tus^−^) strains of *E. coli*. All plasmids used for transfection were prepared by endotoxin-free maxiprep (QIAGEN Sciences). siRNA SMARTpools were purchased from Horizon Scientific/Dharmacon.

### Cell line generation and cell culture

Frt-F13 and related reporter cell lines were generated in conditional mouse *Brca1*^fl/hyg^ ES cell (mES) line. A single copy of the reporter was integrated into the *ROSA26* locus using electroporation. Targeted integration and copy number were verified using Southern blotting, as described previously^26^. *BRCA1*-deficient (*Brca1*^Δ11/hyg^) ES clones were generated by transient adenovirus-mediated Cre expression. mES cells were thawed on irradiated mouse embryonic fibroblast (MEF) feeder cells and maintained on gelatinized plates. Cell lines are routinely tested for mycoplasma infection by Myco-Alert assay (Lonza).

### TrAEL-seq

Mouse ES cells in a 129S6/SvEvTac background were maintained in Dulbeco’s modified eagle medium (Invitrogen) supplemented with 15% FCS (Invitrogen), L-Glutamine (Gibco), NEAs (Gibco), B-mercaptoethanol (Sigma) and LIF and cultured on plates coated with 0.2% gelatin (Sigma). The preparation and bioinformatic analysis of TrAEL-seq libraries followed the method described previously^42^. The data has been deposited in GEO (accession number GSE259364).

### Recombination assays

Before transfection, adherent mES cells are detached using 0.25% Trypsin and quenched using serum. In suspension, 1.6 x 10^5^ cells were transfected with 0.5 μg empty vector, pcDNA3b-myc-NLS-wtFlp, pcDNA3b-myc-NLS-FlpH305L, pcDNA3b-myc-NLS-FlpY343, pcDNA3b-myc-NLS-FlpH305L/Y343F double mutant, pcDNA3b-myc-NLS-Tus or pcDNA3b-myc-NLS-I-SceI plasmid using Lipofectamine 2000 (Invitrogen). Co-transfections for the Frt-F13-6xTer reporter cell line were performed with 0.5 μg Tus and 0.5 μg FlpH305L using Lipofectamine 2000. In these experiments, 0.5 μg Tus and 0.5 μg FlpH305L single transfections were co-transfected with 0.5 μg EV to maintain the same nucleic acid amounts. GFP^+^RFP^−^, GFP^+^RFP^+^ and GFP^−^RFP^+^ repair outcomes were detected 72 hours after transfection and measured using flow cytometry Beckman Coulter CytoFlex LX. Samples were transfected in technical duplicates and 3-6 x 10^5^ total events were scored. Transfection efficiencies were measured for each experiment using 0.05 μg *GFP* expression vector and 0.45 μg EV. All the data presented are corrected for background noise (EV transfection only) and for transfection efficiency (40-90%).

### Analysis of repair outcomes using Southern blotting

Sorted GFP^+^RFP^−^, GFP^+^RFP^+^ and GFP^−^RFP^+^ repair outcomes were cultured and expanded to reach ∼80% confluence. Genomic DNA was isolated from these clones using a Puregene DNA Isolation Kit (QIAGEN Sciences, Maryland, MD). Southern blotting of BglII and BglII + I-SceI digested genomic DNA was performed using a *GFP* cDNA as a radiolabeled probe as described previously^22^.

### Statistical methods

One-way ANOVA analysis was performed for samples greater than three (n>3). Each plotted data point is calculated as a mean of technical duplicates. Each dataset is presented as the arithmetic mean of all the data points and the error bars represent the standard error of the mean (s.e.m.) of ≥ 6 independent experiments. Other experiments where shown were analyzed using Student’s two-tailed unpaired *t*-test, assuming unequal variance. No statistical methods were used to predetermine sample size. The experiments were not randomized, and investigators were not blinded to allocation during experiments and outcome assessment. Statistics were performed using GraphPad Prism v10.1 software.

