## Supplemental Figures 1-7 for "Two-ended recombination at a Flp-nickase-broken replication fork"

**Supplemental Figure Legends**

**Supplemental Figure S1. Analysis of FlpH305L-induced HR products.** Related to **Figure 1**. **A.** TrAEL-seq data in mES cells, showing regions flanking *Rosa26* in Chr 6. Read polarity calculated as described in STAR Methods. Arrow shows transcription direction of *Rosa26*. **B.** FlpH305L binding to the *Frt*-F13min site generates DPC nicks on either strand. Pink: non-incising Flp monomers. Dark red: incising Flp monomer; active site tyrosine (Y) is shown. Lower panel: sequence of *Frt*-F13min site in HR reporter with incision sites. **C.** Western blot showing abundance of Flp products, detected using anti-Myc Ab. β-tubulin: loading control. **D.** FlpH305L-induced STGC in 6 independent *Frt*-F13 HR reporter clones and 5 independent *Frt*-F13min HR reporter clones. Data shows mean and s.e.m. of 6 independent experiments (n=6). Mean of all pooled *Frt*-F13 HR reporter data points = 0.070%. Mean of all pooled *Frt*-F13min HR reporter data points = 0.054%. Unpaired *t*-test of pooled data points for *Frt*-F13 HR reporter clones *vs*. *Frt*-F13min HR reporter clones: P=0.0013. **E.** Representative raw FACS plots of Flp mutant-induced repair products in F13min HR reporter clone, uncorrected for transfection efficiency. Note STGC ($GFP^{+}RFP^{-}$), LTGC ($GFP^{+}RFP^{+}$) and $GFP^{-}RFP^{+}$ outcomes in FlpH305L-transfected cells and absence of induced repair events in Flp Y343F-transfected cells. **F.** Upper panel: Predicted structures of two-ended STGC products and of LTGC products, with BglII and I-SceI restriction fragments shown**.** Lower panel: Southern blot analysis of gDNA from LTGC (L1-4) and STGC (S1-4) clones, digested with BglII (B) or BglII + I-SceI (B+I) and probed with *GFP* cDNA. Note fixed size of STGC products reflecting two-ended HR.

**Supplemental Figure S2. Genetic analysis of FlpH305L-induced and I-SceI-induced repair.** Related to **Figure 2**. **A** and **B.** FlpH305L-induced I-SceI induced (A) and FlpH305L-induced (B) repair in parental *Xrcc4*$^{fl/fl}$ and *Xrcc4*$^{-/-}$ reporter cells. Left panel: LTGC. Middle panel: Ratio of LTGC:Total $GFP^{+}$. Right panel: $GFP^{-}RFP^{+}$ frequencies Data shows mean and s.e.m. of 4 independent experiments (n=4). Unpaired *t*-test: ns: not significant; * P < 0.05; ** P < 0.01. **C-F.** Frequencies of I-SceI-induced repair outcomes in cells depleted of BRCA2 and/or Rad51. Cells were co-transfected with I-SceI and siRNAs as shown. See STAR Methods for experimental details. Data shows mean and s.e.m. of 8 independent experiments (n=8). Unpaired *t*-test: ns: not significant; * P < 0.05; ** P < 0.01. **C.** Left panel: STGC frequencies. Middle and right panels: abundance of *Brca2* and *Rad51* mRNA in siRNA-transfected cells detected by RT-qPCR. **D.** I-SceI-induced LTGC frequencies. **E.** Ratio of LTGC:Total $GFP^{+}$ frequencies. **F.** Frequencies of I-SceI-induced $GFP^{-}RFP^{+}$ outcomes.

**Supplemental Figure S3. Analysis of I-SceI-induced $GFP^{-}RFP^{+}$ products.** Related to **Figure 3**. **A.** Southern blot analysis of 9 independent I-SceI-induced $GFP^{-}RFP^{+}$ products. Clone #3 was lost. Genomic DNA was digested *in vitro* with BglII (B) or with BglII+I-SceI (B+I), *GFP* probe. Note: all clones except #2 and #5 (red *) contain two I-SceI sites, similar to major FlpH305L-induced $GFP^{-}RFP^{+}$ products. **B.** Proposed mechanism of generation of FlpH305L-induced and I-SceI-induced $GFP^{-}RFP^{+}$ outcome *via* discoordinated dual invasions of the sister chromatid. Original sites of left and right DNA ends are marked L and R, respectively.

**Supplemental Figure S4. *Brca1* exon 11 is dispensable for FlpH305L-induced STGC.** Related to **Figure 4**. **A**. PCR primers used to identify *Brca1*$^{fl}$ and *Brca1*$^{\Delta 11}$ clones. Primers *a* and *b* identify *Brca1*$^{fl}$; primers *a* and *c* identify *Brca1*$^{\Delta 11}$. **B-E**. Data from 6 independent Cre-exposed Brca1$^{fl/hyg}$ clones and 6 independent Cre-exposed Brca1$^{\Delta 11/hyg}$ clones, from which pooled data in **Figure 4** is derived. Data shows mean and s.e.m. of 6 independent experiments (n=6). **B.** STGC. **C.** LTGC. Note elevated FlpH305L-induced LTGCs in Brca1$^{\Delta 11/hyg}$ cells. **D.** Ratio of LTGC:Total GFP$^{+}$. **E.** FlpH305L-induced GFP$^{-}$RFP$^{+}$ outcomes.

**Supplemental Figure S5. Domains of BRCA1 additional to exon 11 control FlpH305L-induced STGC.** Related to **Figure 4E**. Brca1$^{fl/hyg}$ clone #12 and Brca1$^{\Delta 11/hyg}$ clone #8 (see **Supplemental Figure S4**) were co-transfected with siRNAs shown together with either FlpH305L or I-SceI, as described in STAR methods. Data from same experiments as in **Figure 4E**. Data shows mean and s.e.m. of 4 independent experiments (n=4). Unpaired *t*-test: ns: not significant; * $P < 0.05$; ** $P < 0.01$; *** $P < 0.001$; **** $P < 0.0001$. Key below data shows the color code for siRNA and FlpH305L or I-SceI transfections. **A.** LTGC. **B.** Ratio of LTGC:Total GFP$^{+}$ frequencies. **C.** Frequencies of FlpH305L-induced GFP$^{-}$RFP$^{+}$ outcomes.

**Supplemental Figure S6. FlpH305L-induced STGC is the product of a single fork collision with the Flp-nick.** Related to **Figure 5. A.** CUT&RUN qPCR for HA-Tus in regions immediately adjacent to the *Ter* array. Empty vector controls normalized to 1.0. See STAR Methods for experimental details. **B.** CUT&RUN qPCR for Rad51 in regions immediately adjacent to the *Ter* array. Empty vector controls normalized to 1.0. See STAR Methods for experimental details. **C-F.** Data from 5 independent *Brca1*$^{fl/hyg}$ clones targeted with the F13-HR reporter carrying an additional 6x*Ter* array positioned to establish Tus-mediated barrier to the leftward fork, from which pooled data in **Figure 5** is derived. Data shows mean and s.e.m. of 5 independent experiments (n=5). **C.** STGC. **D.** LTGC. **E.** Ratio of LTGC:Total $GFP^{+}$. **F.** FlpH305L-induced $GFP^{-}RFP^{+}$ outcomes. **G.** Southern blot analysis of FlpH305L-induced LTGC products in presence of Tus/*Ter* RFB. Genomic DNA was digested with BglII, *GFP* probe. M: Molecular weight markers. **H.** Southern blot analysis of FlpH305L-induced $GFP^{-}RFP^{+}$ products in presence of Tus/*Ter* RFB. Genomic DNA was digested with BglII, *GFP* probe. M: Molecular weight markers.

**Supplemental Figure S7. Model: collision of a single replication fork with a leading strand Flp-nick generates a one-ended break.** Related to **Figure 6.** A Flp-nick on the leading strand exposes a free 5' end of the leading parental strand, leading to loss of CMG from the parental strand and termination of DNA synthesis.

Fig_S1

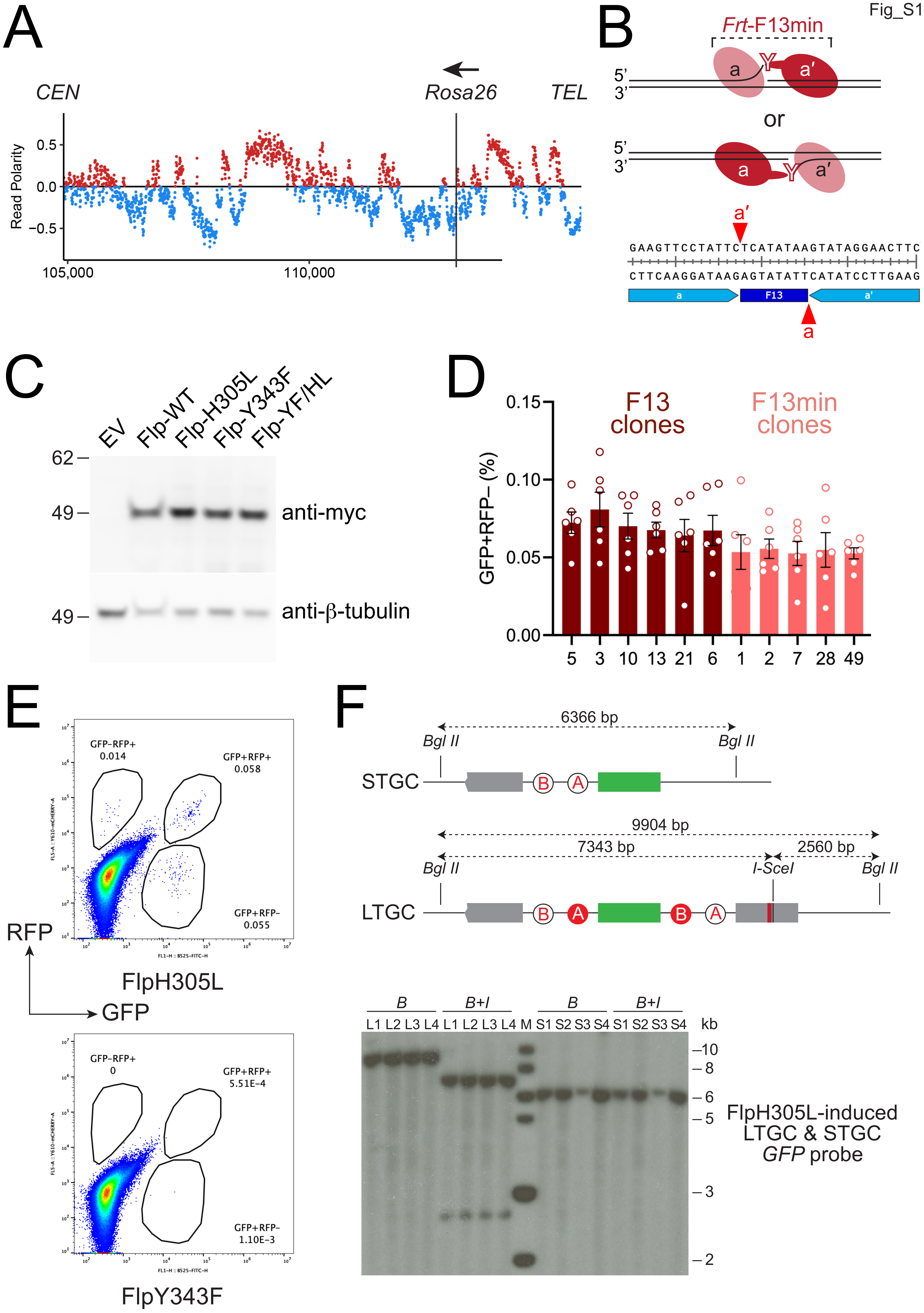

A
CEN
Rosa26
TEL
Read Polarity
0.5
0.0
−0.5
105,000
110,000
B
Frt-F13min
5'
3'
a
a'
or
GAAGTTCCTATTCTCATATAAGTATAGGAACTTC
CTTCAAGGATAAGAGTATATTCATATCCTTGAAG
F13
C
EV
Flp-WT
Flp-H305L
Flp-Y343F
Flp-YF/HL
62
49
anti-myc
anti-β-tubulin
D
F13 clones
F13min clones
GFP+RFP− (%)
0.15
0.10
0.05
0.00
5
3
10
13
21
6
1
2
7
28
49
E
GFP−RFP+
0.014
GFP+RFP+
0.058
GFP+RFP−
0.055
RFP
GFP
FlpH305L
GFP−RFP+
0
GFP+RFP+
5.51E-4
GFP+RFP−
1.10E-3
FlpY343F
F
6366 bp
Bgl II
STGC
B
A
9904 bp
7343 bp
2560 bp
I-SceI
LTGC
B
B+I
L1 L2 L3 L4 L1 L2 L3 L4 M S1 S2 S3 S4 S1 S2 S3 S4
kb
10
8
6
5
3
2
FlpH305L-induced
LTGC & STGC
GFP probe

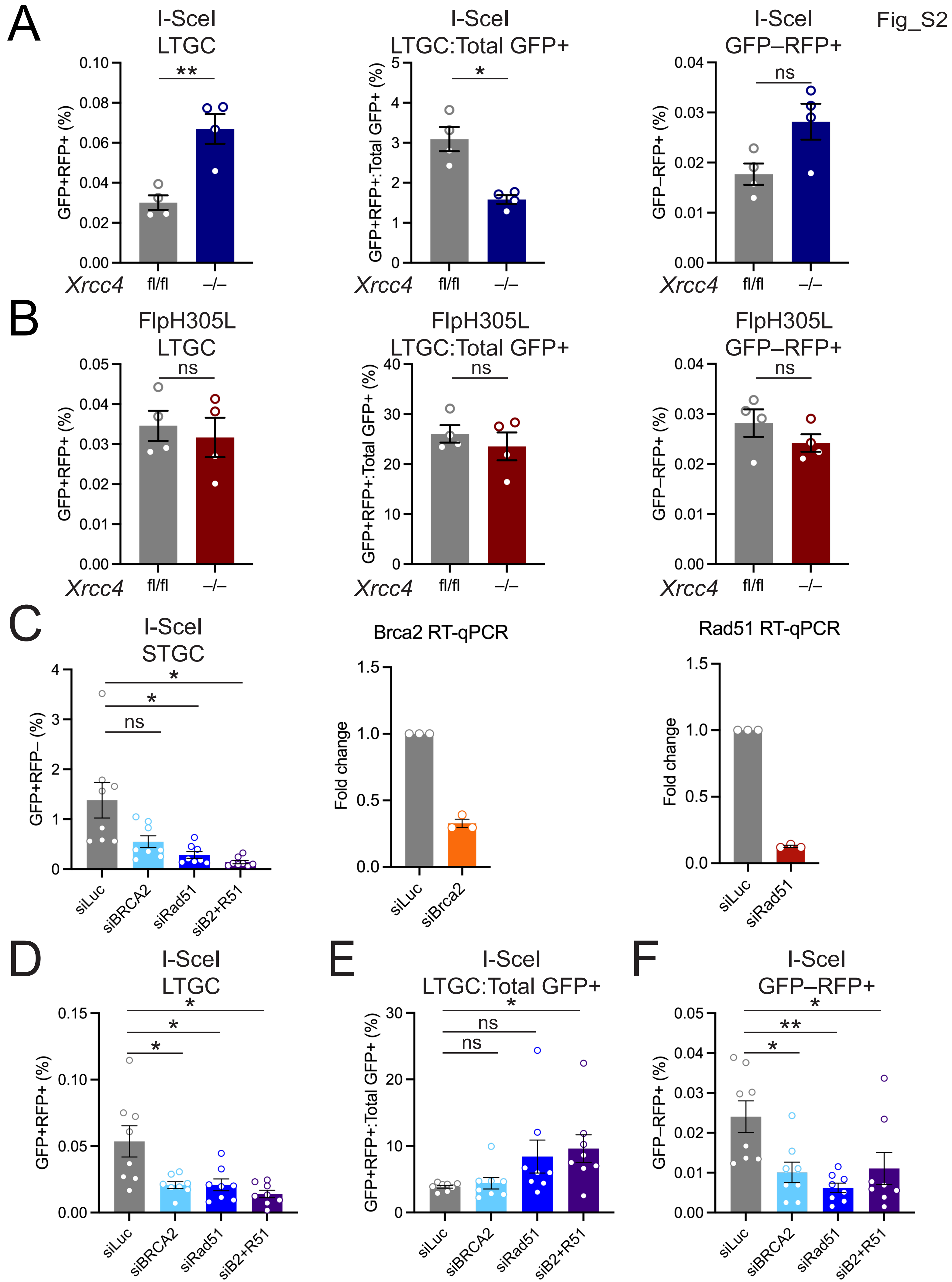

Fig_S2
A
I-SceI
LTGC
GFP+RFP+ (%)
**
I-SceI
LTGC:Total GFP+
GFP+RFP+:Total GFP+ (%)
*
I-SceI
GFP–RFP+
GFP–RFP+ (%)
ns
Xrcc4
fl/fl
–/–
B
FlpH305L
LTGC
FlpH305L
LTGC:Total GFP+
FlpH305L
GFP–RFP+
C
I-SceI
STGC
GFP+RFP– (%)
siLuc
siBRCA2
siRad51
siB2+R51
Brca2 RT-qPCR
Fold change
siBrca2
Rad51 RT-qPCR
D
I-SceI
LTGC
E
I-SceI
LTGC:Total GFP+
F
I-SceI
GFP–RFP+

Fig_S3

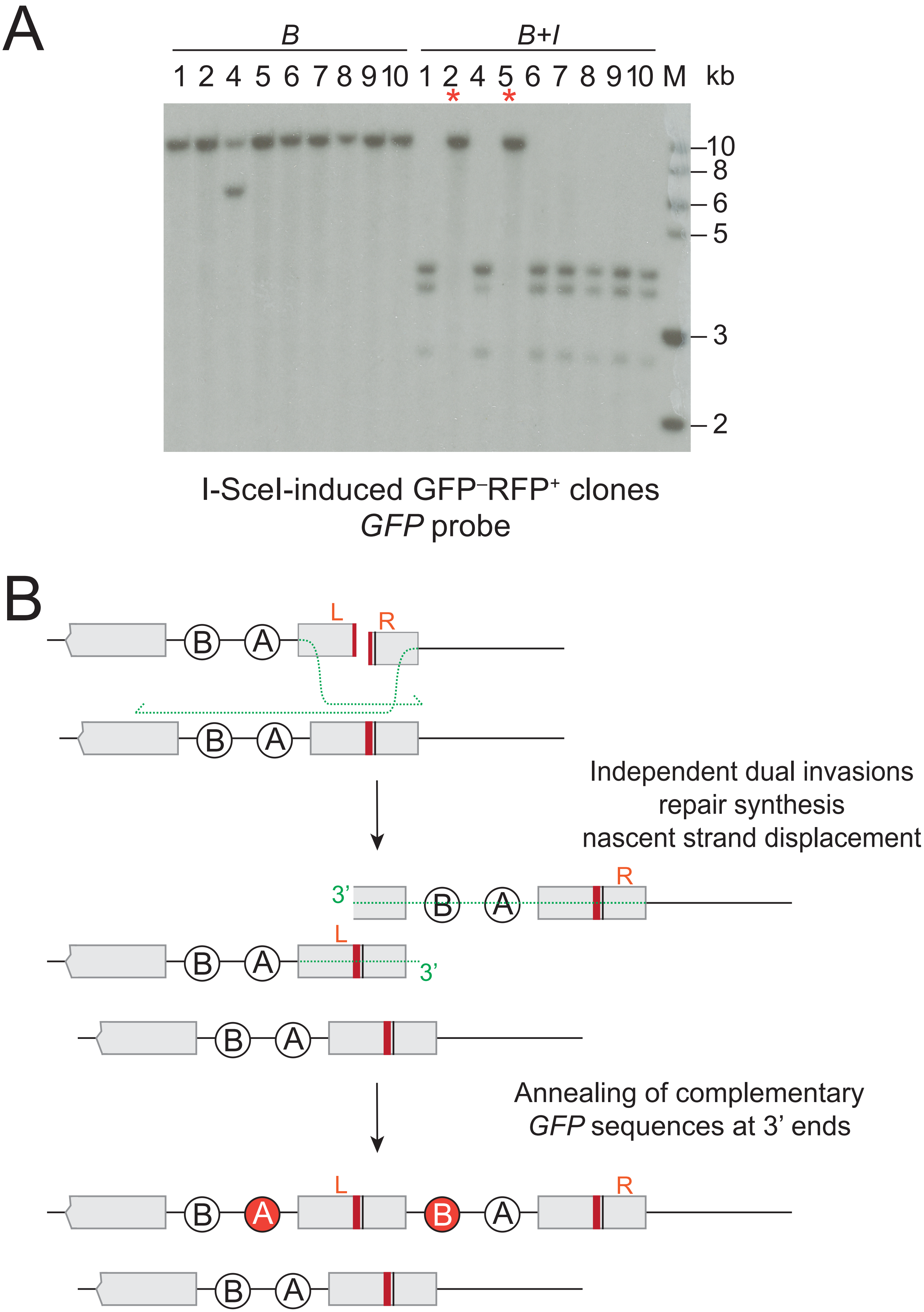
A
B
B+I
1 2 4 5 6 7 8 9 10 1 2 4 5 6 7 8 9 10 M kb
10
8
6
5
3
2
I-SceI-induced GFP⁻RFP⁺ clones
GFP probe
B
L
R
B
A
Independent dual invasions
repair synthesis
nascent strand displacement
3'
Annealing of complementary
GFP sequences at 3' ends

Fig_S4

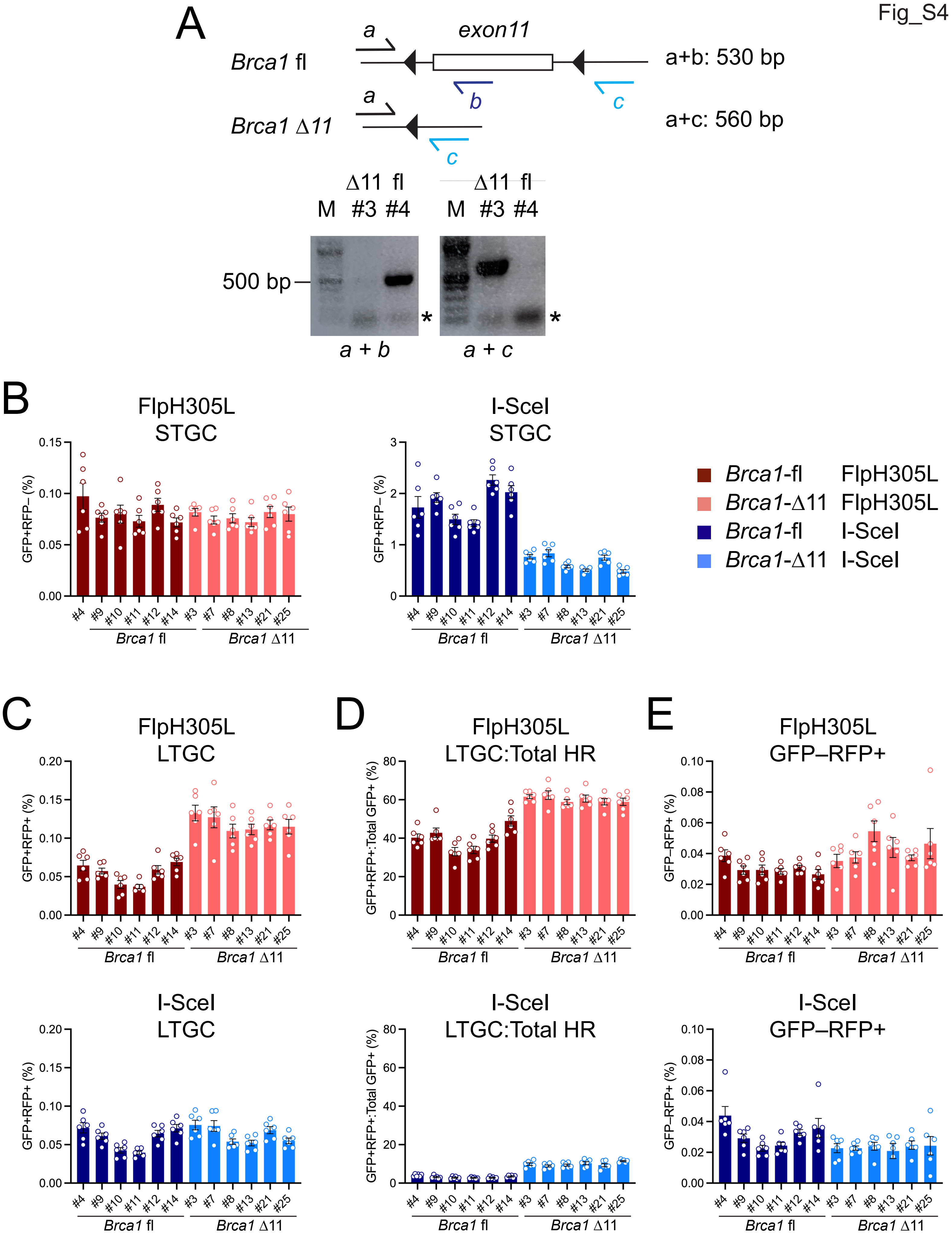
A
Brca1 fl
a
exon11
b
c
a+b: 530 bp
Brca1 Δ11
a
c
a+c: 560 bp
Δ11 fl
M #3 #4
Δ11 fl
M #3 #4
500 bp
*
*
a + b
a + c
B
FlpH305L STGC
I-SceI STGC
GFP+RFP– (%)
Brca1 fl
Brca1 Δ11
Brca1-fl FlpH305L
Brca1-Δ11 FlpH305L
Brca1-fl I-SceI
Brca1-Δ11 I-SceI
C
FlpH305L LTGC
GFP+RFP+ (%)
D
FlpH305L LTGC:Total HR
GFP+RFP+:Total GFP+ (%)
E
FlpH305L GFP–RFP+
GFP–RFP+ (%)
I-SceI LTGC
I-SceI LTGC:Total HR
I-SceI GFP–RFP+
#4 #9 #10 #11 #12 #14 #3 #7 #8 #13 #21 #25

Fig_S5

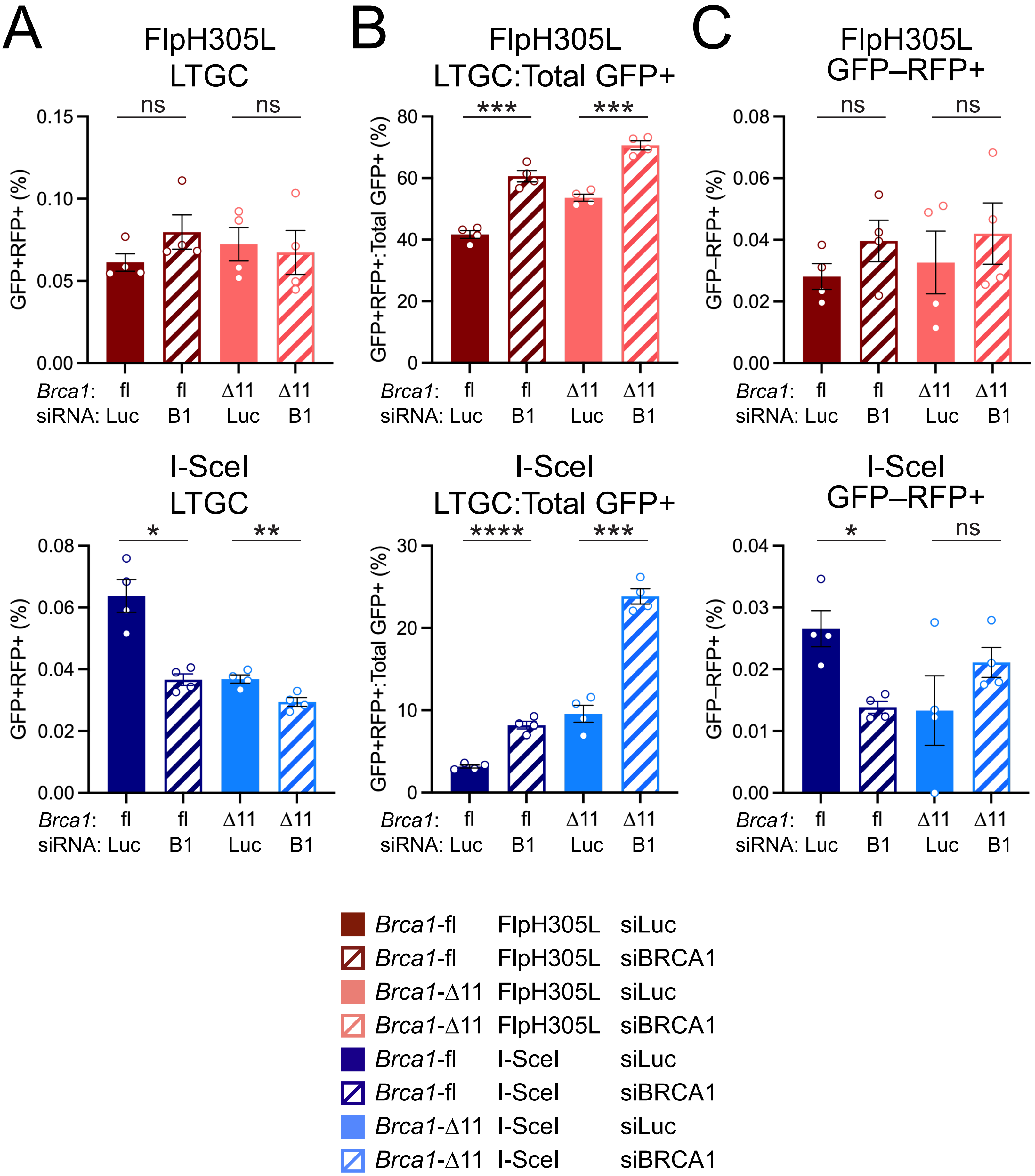
A
FlpH305L
LTGC
GFP+RFP+ (%)
0.15
0.10
0.05
0.00
ns
ns
Brca1: fl fl Δ11 Δ11
siRNA: Luc B1 Luc B1
B
FlpH305L
LTGC:Total GFP+
GFP+RFP+:Total GFP+ (%)
80
60
40
20
0
***
***
Brca1: fl fl Δ11 Δ11
siRNA: Luc B1 Luc B1
C
FlpH305L
GFP–RFP+
GFP–RFP+ (%)
0.08
0.06
0.04
0.02
0.00
ns
ns
Brca1: fl fl Δ11 Δ11
siRNA: Luc B1 Luc B1
I-SceI
LTGC
GFP+RFP+ (%)
0.08
0.06
0.04
0.02
0.00
*
**
Brca1: fl fl Δ11 Δ11
siRNA: Luc B1 Luc B1
I-SceI
LTGC:Total GFP+
GFP+RFP+:Total GFP+ (%)
30
20
10
0
****
***
Brca1: fl fl Δ11 Δ11
siRNA: Luc B1 Luc B1
I-SceI
GFP–RFP+
GFP–RFP+ (%)
0.04
0.03
0.02
0.01
0.00
*
ns
Brca1: fl fl Δ11 Δ11
siRNA: Luc B1 Luc B1
Brca1-fl FlpH305L siLuc
Brca1-fl FlpH305L siBRCA1
Brca1-Δ11 FlpH305L siLuc
Brca1-Δ11 FlpH305L siBRCA1
Brca1-fl I-SceI siLuc
Brca1-fl I-SceI siBRCA1
Brca1-Δ11 I-SceI siLuc
Brca1-Δ11 I-SceI siBRCA1

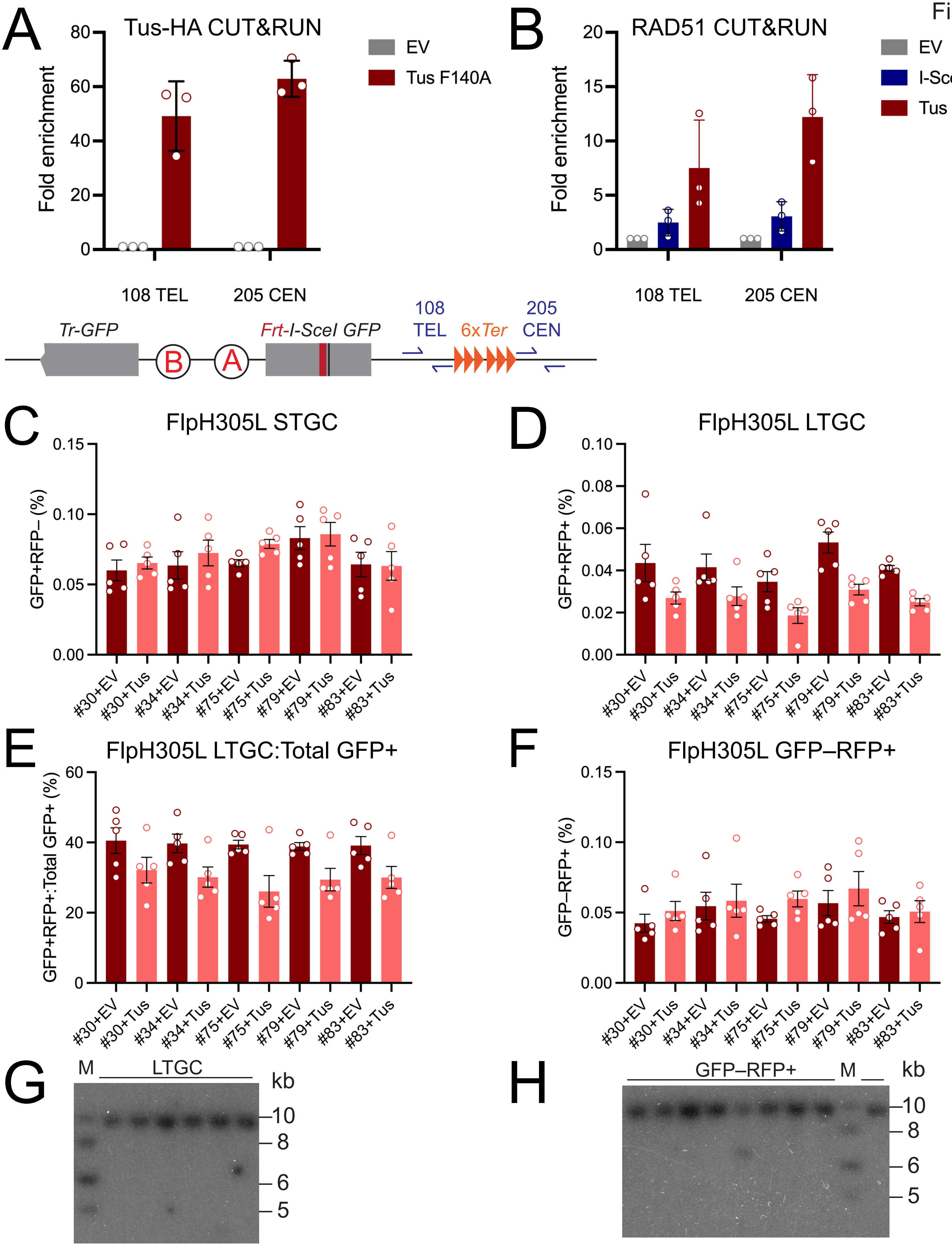


FlpH305L + Tus
LTGC clones
Bgl II digest, *GFP* probe

FlpH305L + Tus
GFP–RFP+ clones
Bgl II digest, *GFP* probe

Fig_S7

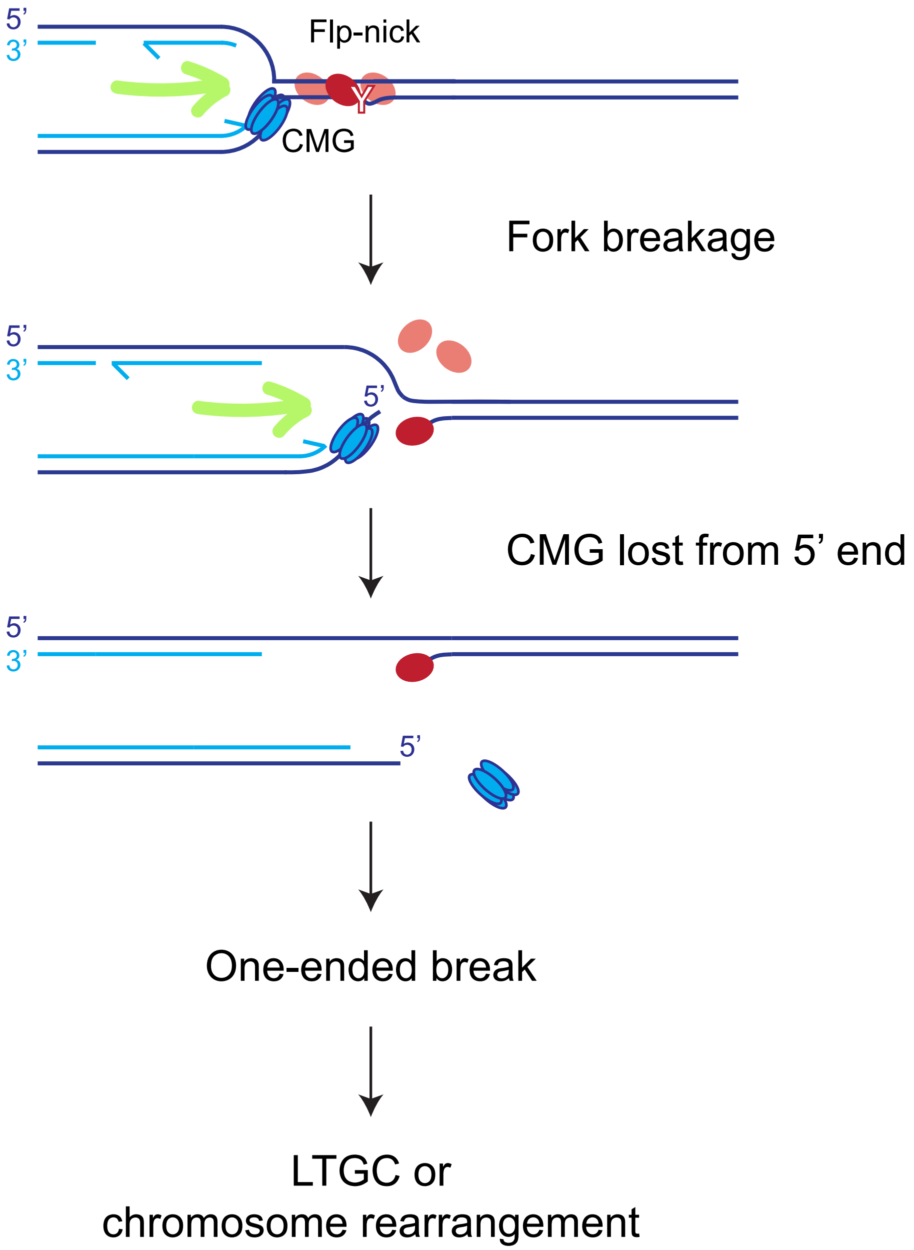

5'
3'
Flp-nick
CMG
Fork breakage
5'
3'
5'
CMG lost from 5' end
5'
3'
5'
One-ended break
LTGC or
chromosome rearrangement
